# Pannexin 1 phosphorylation sites differentially modulate channel activity and physiological outcomes

**DOI:** 10.64898/2026.07.12.738047

**Authors:** Brooke L. O’Donnell, Luke S. Dunaway, Xuexin Zhang, Skylar A. Loeb, Zuzanna J. Juśkiewicz, Wyatt J. Schug, Susan A. Leonhardt, Abigail G. Wolpe, Melissa A. Luse, Samantha C. Bielefeld, Andrew K.J. Boyce, Madison D. Williams, Marie Billaud, Angela K. Best, Scott R. Johnstone, Silvia Penuela, Linda Columbus, Anastasia F. Thévenin, Roger J. Thompson, Douglas A. Bayliss, Michael Koval, Brant E. Isakson

## Abstract

Within the vasculature, pannexin 1 (PANX1) channels in smooth muscle cells (SMCs) regulate α-adrenergic constriction and blood pressure. PANX1 channel activity is regulated by phosphorylation at Y198, S205 and Y308 residues, but the physiological significance of these modifications is unknown. Here, we utilize newly developed PANX1 Y198F, S205A and Y308F phospho-dead mutant mice to test physiological changes related to hemodynamics. Radiotelemetry-measured blood pressure was decreased in Y198F, increased in Y308F, but unchanged in S205A mice at baseline. Clonidine-sensitive sympathetic-driven hypertension was observed in all mouse lines except Y198F. Pressure myography of third-order mesenteric arteries revealed α-adrenergic contractile responses were decreased in Y198F, slightly enhanced in Y308F, but unchanged in S205A, with responses in Y198F vessels mimicking controls treated with PANX1 inhibitors. To understand signaling changes driving these phenotypes, we performed mesenteric artery bulk RNA sequencing, but found a minimal number of differentially expressed genes between phospho-dead mutants and controls. Similarly, co-immunoprecipitation-mass spectrometry of wildtype or phospho-dead mutant-expressing vascular SMCs revealed few interacting proteins distinct to each PANX1 variant. However, PANX1 channel activity assessments in HEK293T cells expressing the α1D-adrenergic receptor as well as each phospho-dead mutant PANX1 showed that phenylephrine-induced ATP release from Y198F channels was significantly decreased compared to wildtype, but current was unaffected. Conversely, basal and phenylephrine-induced S205A and Y308F currents were reduced, but ATP release resembled controls. Taken together, these findings indicate that distinct PANX1 phosphorylation determines PANX1 metabolite release versus current conducting properties and in turn, regulates physiological outcomes in the vasculature.

**One Sentence Summary:** PANX1 Y198 phosphorylation-mediated ATP release is a major driver of α-adrenergic vasoconstriction in vascular smooth muscle cells.

## INTRODUCTION

Hypertension is extremely common among adults worldwide and is a major risk factor for cardiovascular disease (*1–3*). Almost 20% of hypertensive patients have treatment-resistant hypertension, where blood pressure is uncontrolled despite the use of ≥3 anti-hypertensives, and the risk of heart failure, end stage renal disease and death is significantly increased (*2, 4, 5*). Due to its high prevalence and treatment burden (*6, 7*), it is critical to understand factors that contribute to the maintenance of healthy blood pressure as well as the development of hypertension.

Pannexin 1 (PANX1) is a large-pore heptameric channel (*8–10*) and scaffold protein (*11–13*) involved in cellular communication that is present in the vascular smooth muscle cells (SMCs) of resistance arteries which regulate blood pressure (*14*). Upon α-1D adrenergic receptor (α-1DAR) activation by naturally occurring and synthetic agonists (norepinephrine and phenylephrine, respectively) or sympathetic nerve activity, α-1DAR receptor signaling activates cell surface PANX1 and facilitates ATP release to drive vascular processes such as vasoconstriction and blood pressure (*15–19*). Resistance artery contraction and ATP release stimulated by α-adrenergic signaling is specific to vascular SMC PANX1 since both processes could be decreased by SMC-specific genetic deletion of *Panx1* (*16*), and increased by overexpression of human PANX1 in SMCs (*19*). By contrast, these measures were unchanged in endothelial cell (EC)-specific *Panx1* knockout (KO) vessels compared to controls (*20*). Furthermore, vasoconstriction through α-adrenergic receptor stimulation was shown to be dependent on ATP, PANX1 and metabotropic purinergic receptor (P2YR) signaling since apyrase, PANX1 channel blockers and purinergic inhibitors abolished the contractile effects (*18–20*). Consistent with these findings, SMC-specific *Panx1* KO mice have reduced nocturnal blood pressure (*16*), whereas mice with human PANX1 overexpression (iSMC PANX1 OE) exhibit elevated baseline blood pressure that is inhibited acutely with injection of the PANX1 inhibitor spironolactone (SPIR) (*19*). Again, EC-specific *Panx1* KO mice did not show any differences in blood pressure compared to controls (*21*). Building on these studies using mouse models, we previously observed Y198-phosphorylated PANX1 (pY198) was upregulated in vascular SMCs of gluteal arteries from hypertensive patients compared to normotensive controls (*17*). Additionally, PANX1 channel inhibition by SPIR or PxIL2P peptide (which targets the intracellular loop region of PANX1 containing Y198) markedly inhibited phenylephrine-induced contractile responses of human arterioles isolated from treatment resistant hypertension patients (*20*). These findings suggest that phosphorylation of PANX1 at Y198 could contribute to blood pressure homeostasis, but it is unknown how changes to PANX1 biochemistry could promote organism level conditions such as hypertension.

In a recent breakthrough study on related connexin hemichannels, Gaete *et al.* discovered that small molecule transport can be uncoupled from ion conductance, and that these processes can be differentially regulated. Moreover, previous classifications of connexin mutations as ‘loss of function’ may not be entirely accurate, since one aspect of channel activity may be altered whereas the other is unaffected (*22*). The recent surge of cryogenic electron microscopy (cryo-EM) structures of PANX1 has suggested that these channels can support both small molecule and ion movement independently through different channel compartments (*10*). In these models, the central pore of PANX1 facilitates the permeation of small molecules less than 1 kDa in size, like ATP as well as ions (*8–10, 23–28*), whereas the lateral/side tunnels located between the heptamer subunits conduct ions such as Cl^−^ (*10, 28*). PANX1 structural studies have also discussed the importance of the N- and C-termini in gating ion and small molecule movement through the channel—with some conflicting evidence depending on cryo-EM sample preparation and the PANX1 variant used (*8–10, 23–27*)—but the possible role of post-translational modifications in supporting these distinct channel functions is yet to be investigated. We hypothesized that PANX1 phosphorylation differentially regulates ATP release and ion conductance, and that these distinct channel properties underlie divergent physiological outcomes.

Post-translational modifications like glycosylation, S-nitrosation and phosphorylation influence PANX1 characteristics including localization and channel activity (*17, 29–33*). In particular, phosphorylation on tyrosine (Y) and serine (S) mouse PANX1 residues Y198, S205 and Y308 (Y199, S206 and Y309, respectively, in human PANX1) located in either the intracellular loop or cytoplasmic C-terminal tail modulate PANX1 channel activity (Fig. 1A). We have previously shown that PANX1 ATP release stimulated by α-adrenergic receptor agonist phenylephrine is supported by constitutive Src-mediated Y198 phosphorylation of PANX1 in arterial SMCs (*17*), whereas in venous ECs, PANX1-mediated ATP release is induced by tumor necrosis factor α (*29*). In HEK293T cells, salt-inducible kinase phosphorylation of S205 stimulates PANX1 currents downstream of α-adrenergic signaling, and a PANX1 S205A mouse line (with the inability of PANX1 to be phosphorylated at the S205 site globally) was used to show that the salt-inducible kinase-PANX1 axis in T cells is critical in reducing airway inflammation in an asthma model (*34*). In cardiomyocytes, β1-adrenergic or histamine-H2-receptor signaling triggered protein kinase A phosphorylation of PANX1 S206, promoting PANX1-mediated ATP release and ultimately resulting in cardiomyocyte cell death (*35*). Conversely, nitric oxide-induced protein kinase G phosphorylation of PANX1 in HEK293 cells, which was postulated to occur on residue S206, attenuated PANX1 ionic current (*32*). Y308 phosphorylation of PANX1 downstream of N-methyl-D-aspartate receptor (NMDAR) signaling in neurons induced PANX1-mediated ion flux based on studies using the TAT-Panx_308 p_eptide which prevents Src interaction with PANX1 at the Y308 phosphorylation site (*33, 36*). Furthermore, during hippocampal NMDAR long-term synaptic depression, non-ionotropic NMDAR signaling may stimulate PANX1-mediated ATP release by Src-dependent Y308 phosphorylation (*37*). Collectively, these results demonstrate the diversity of signals which converge on PANX1 in different cell types, as well as the possibility that molecular versus ion permeation through the PANX1 channel may be distinctly modified depending on the PANX1 phosphosite. However, despite these findings on the importance of phosphorylation in the regulation of PANX1 channel function, the physiological consequences of PANX1 pY198, pS205 and pY308 are not known, nor are the molecular mechanisms linking these post-translational modifications to vascular physiology.

**Fig 1.**
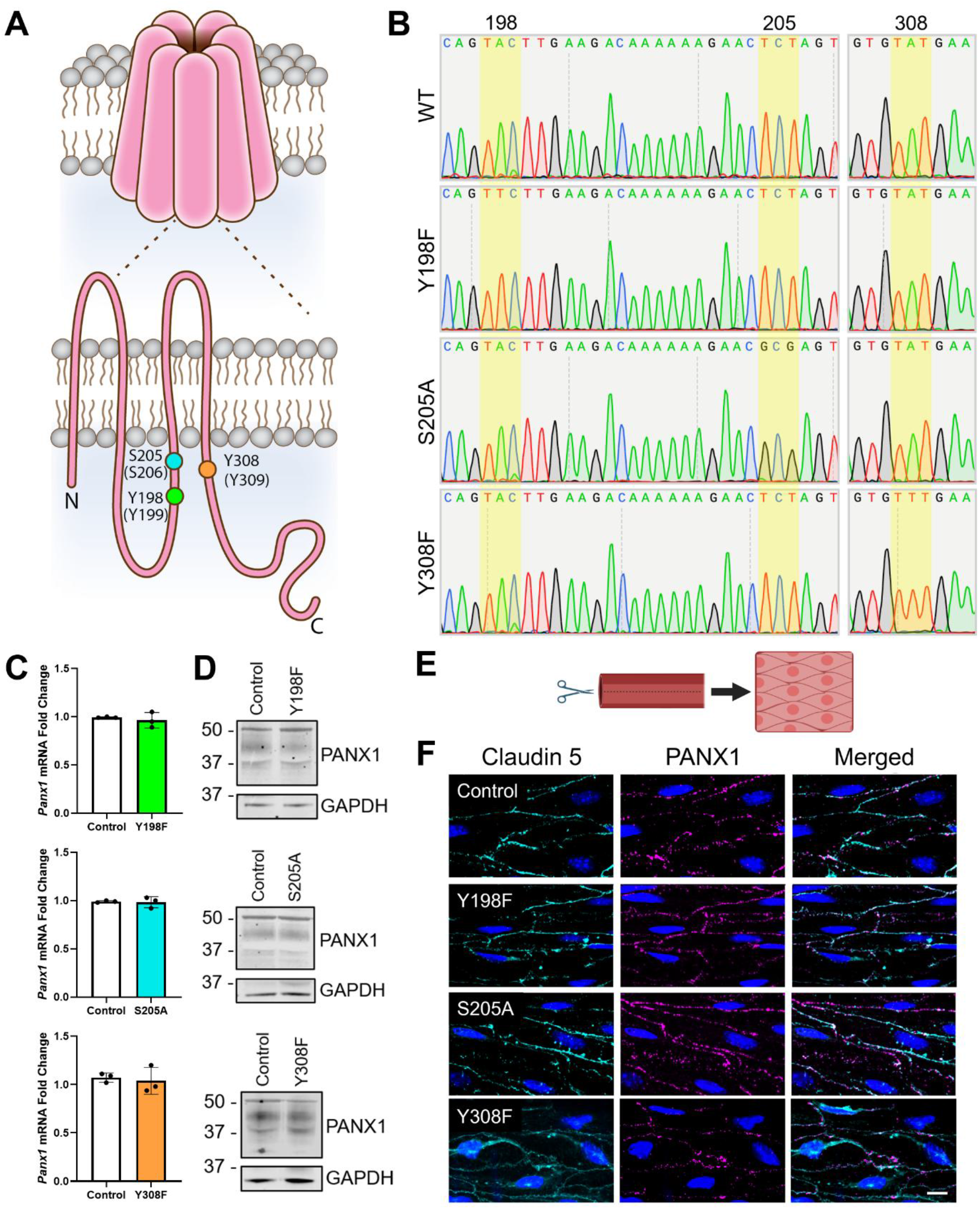
PANX1 Y198F, S205A and Y308F phospho-dead mutants exhibit normal expression and trafficking. (**A**) Schematic of PANX1 oligomerization and topology with the location of Y198 (green), S205 (blue) and Y308 (orange) mouse PANX1 phosphosites. Corresponding human PANX1 residues in brackets. N-terminal and C-terminal tails denoted by N and C, respectively. (**B**) Sanger sequencing of tail snips from wildtype control (WT) or global Y198F, S205A and Y308F phospho-dead mutant mice confirmed missense mutations for each mouse line. Genomic DNA corresponding to mouse PANX1 residues 198, 205 and 308 outlined in yellow. (**C**) RT-qPCR of *Panx1* levels in mesenteric arteries from phosphomutant mice and littermate controls. Unpaired t-test. *N*=3 mice. Bars represent mean ± SD. (**D**) Immunoblotting of mesenteric arteries from phosphomutant mice and littermate controls. Protein sizes in kDa. *N*=1 mouse. (**E**) Third-order mesenteric arteries were prepared *en face*, where the vessel is dissected longitudinally and opened to visualize the innermost endothelial cells (schematic created with BioRender.com). (**F**) Endothelial cells were immunostained *en face* for the endothelial cell surface marker claudin 5 (cyan) and PANX1 (magenta). Scale bar 10 µm. *N*=1 mouse.

In this study, we used global PANX1 Y198F, S205A and Y308F phospho-dead mutant mouse models in comparison to EC- and SMC-specific *Panx1* KO mice to explore how these PANX1 phosphorylation sites affect changes related to hemodynamics. Our results provide insight into the differential regulation of molecular permeation versus ion conductance through PANX1 channels, as well as evidence for how local molecular changes to PANX1 can have broad impacts on organismal physiology.

## RESULTS

### PANX1 expression and localization in PANX1 Y198F, S205A and Y308F mice resembles controls

To assess the physiological effects of PANX1 phosphorylation, we generated global PANX1 Y198F and Y308F mice by clustered regularly interspaced short palindromic Repeats (CRISPR)/Cas9 gene editing and utilized the previously developed PANX1 S205A mouse model (*34*). We genotyped the mice by Sanger sequencing, confirming that each phospho-dead mutant mouse model had the specific missense mutation at the intended phosphosites (Fig. 1B, S1). Both *Panx1* mRNA expression (Fig. 1C) and PANX1 protein levels (Fig. 1D) from isolated mesenteric arteries were similar between Y198F, S205A and Y308F mice and their littermate controls, whereas *Panx1* mRNA was significantly reduced in mesenteric arteries from EC-specific (Fig. S2A) and SMC-specific (Fig. S2B) *Panx1* KO mice. Furthermore, *en face* immunofluorescence of isolated third-order mesenteric arteries showed PANX1 Y198F, S205A and Y308F localize to the plasma membrane of ECs, as evidenced by their co-localization with EC cell surface marker claudin 5, resembling wildtype (WT) PANX1 (Fig. 1E,F). Thus, phospho-dead mutant mice exhibit normal PANX1 expression and trafficking.

To further characterize each murine model, we evaluated the size, body composition and litter genotypes from heterozygous crosses of each phospho-dead mutant mouse line. Heterozygous crosses of each mouse line produced litters with genotypes that were not significantly different from expected Mendelian ratios (Fig. S3A). In terms of body size, Y198F and S205A mice resembled littermate controls at 13 weeks of age (Fig. S3B), and EchoMRI tests revealed no changes in total body mass, or fat, lean and water mass normalized to total body mass (Fig. S3C-D). However, Y308F mice were larger than controls (Fig. S3B), and their increased total body mass was driven by an enhanced percent fat mass (Fig. S3E). There was also a nonsignificant trend to a reduction in percent lean mass, but no changes in percent water mass for Y308F mice. No other overt phenotypes were observed in any phospho-dead mutant mouse line.

### Y198F and Y308F mice exhibit opposite blood pressure phenotypes

ATP released from SMC PANX1 channels has been implicated in the sympathetic control of blood pressure (*16, 19*). To investigate whether PANX1 phosphorylation contributes to blood pressure regulation, we performed radiotelemetry (Fig. 2A) on awake, freely moving 11-week-old mice at baseline to measure real-time blood pressure and heart rate (Fig. 2B-E). We confirmed previous findings from our group which established that SMC-specific *Panx1* KO mice, but not EC-specific *Panx1* KO mice, have reductions in baseline mean arterial blood pressure (MAP) with no changes in heart rate (Fig. S2C-D) (*16, 19, 29, 38–40*). Y198F mice had significantly lower MAP during the day and night (Fig. 2C), with more pronounced differences during their active phase at night when murine α-adrenergic receptor activity is the highest (*41*). Conversely, Y308F mice showed the opposite phenotype, where MAP was significantly increased during both time periods (Fig. 2E). There were no differences observed in MAP of S205A mice (Fig. 2D), nor heart rate for any of the phospho-dead mutant mice compared to controls (Fig. 2C-E). These results indicate blood pressure is differentially regulated by these PANX1 phosphorylation sites.

**Fig 2.**
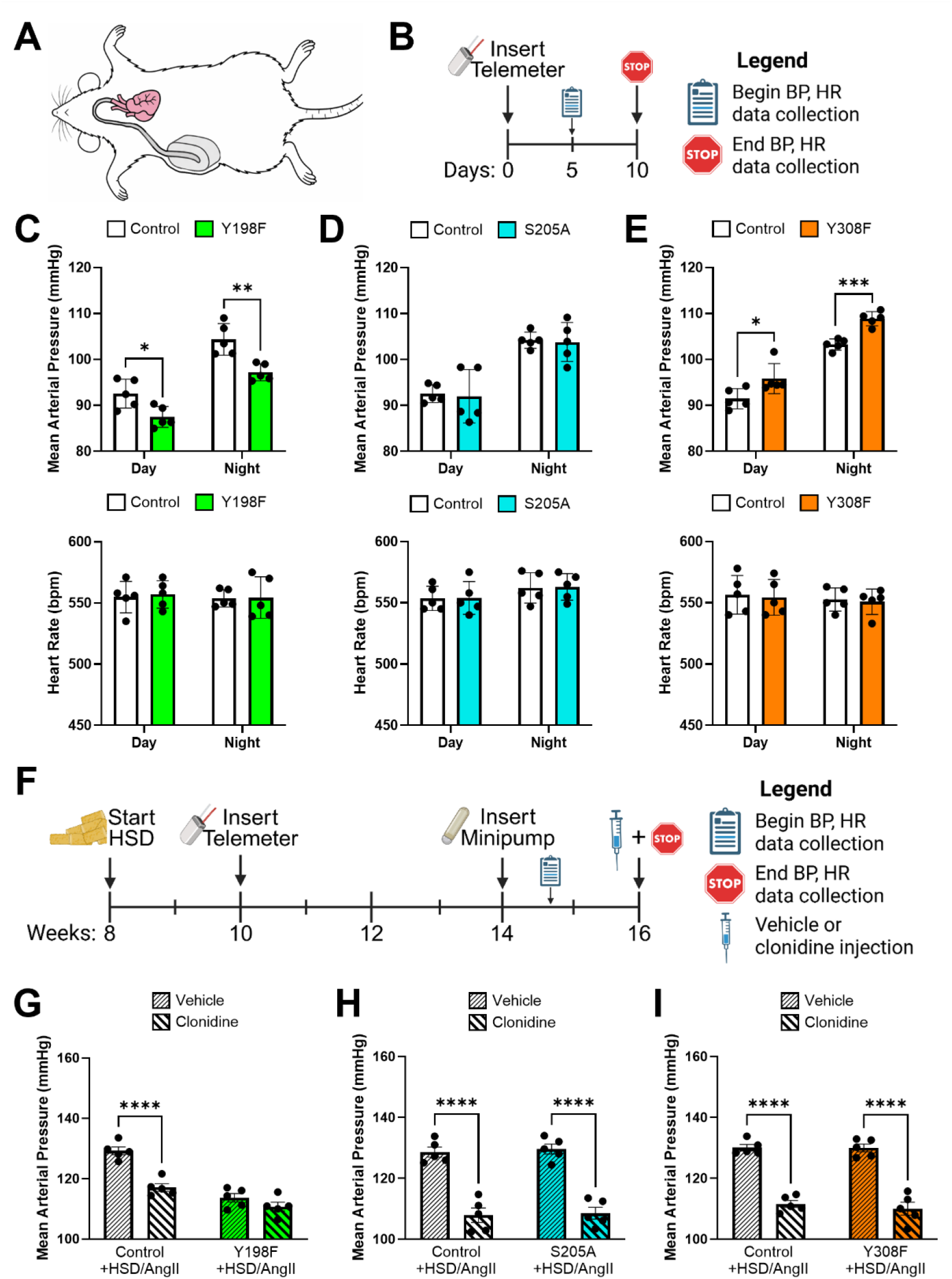
Blood pressure responses of PANX1 Y198F mice are sympathetically driven. (**A**) Mouse schematic showing radiotelemeter insertion into the flank and connection to the carotid artery. (**B**) Experimental timeline for baseline blood pressure (BP) and heart rate (HR) analysis. Radiotelemetry-measured baseline mean arterial pressure and HR during the day and night for Y198F (**C**), S205A (**D**) and Y308F (**E**) mice compared to controls. *N*=5 mice. Unpaired t-test within each time point. \**P*<0.05, \*\**P*<0.01, \*\*\**P*<0.001. (**F**) Experimental timeline for 6% high salt diet (HSD) and 1.0 mg/kg/day angiotensin II (AngII) minipump hypertension model for BP and HR measurement. At 16 weeks of age, mice were injected with either clonidine (0.05 mg/kg) or saline vehicle control. Mean arterial pressure for HSD/AngII Y198F (**G**), S205A (**H**), and Y308F (**I**) mice compared to controls, with and without clonidine treatment. Two-way ANOVA followed by Tukey’s multiple comparisons test. *N*=5 mice, \*\*\*\**P*<0.0001. Bars represent mean ± SD. Schematics created with BioRender.com.

To examine the contribution of PANX1 to sympathetic-driven hypertension, we challenged our mice with a high salt diet and angiotensin II infusion (HSD/AngII) to increase daytime blood pressure (*42*) and then treated with either the sympathetic inhibitor clonidine (*43*) or vehicle control (Fig. 2F). As expected, MAP was elevated with HSD/AngII in control mice (compared to baseline measures in Fig. 2C) and significantly reduced with clonidine compared to vehicle control (Fig. 2G-I). Strikingly, Y198F mice MAP did not increase as much as in control mice after HSD/AngII treatment and was not different from clonidine-treated HSD/AngII control mice with or without clonidine administration. However, MAP of both S205A and Y308F HSD/AngII mice with and without clonidine treatment did not differ significantly from corresponding controls. Taken together, these findings suggest that PANX1 Y198 phosphorylation is a major contributor to blood pressure driven by sympathetic activity.

### Y198F resistance arteries exhibit reduced phenylephrine contractile responses which are unaffected by PxIL2P and SPIR treatment

Blood pressure can be regulated by both cardiac output and total peripheral resistance, with resistance arteries accounting for up to 50% of total peripheral resistance (*44*). Since PANX1 has previously been implicated in α-adrenergic vasoconstriction (*16, 19, 20*), we assessed the contractile responses of isolated third-order mesenteric resistance arteries by pressure myography (Fig. 3A, representative traces Fig. S4) to further understand the contribution of PANX1 phosphorylation to blood pressure regulation. Consistent with published work (*16, 19, 20, 38*), EC-specific *Panx1* KO arteries showed no significant differences in contractile responses to phenylephrine (Fig. S2E), whereas SMC-specific *Panx1* KO vessels had compromised phenylephrine contractile responses (Fig. S2F). This reiterates that PANX1 in SMCs, but not ECs, is critical for α-adrenergic-mediated vessel constriction. EC-specific *Panx1* KO phenylephrine responses could be reduced by treatment with the PxIL2P peptide which targets the Y198 phosphorylation site (*16*) or the PANX1 channel blocker SPIR (Fig. S1G). As expected, treatment with PxIL2P or SPIR did not have any effect on SMC-specific *Panx1* KO vessel response to phenylephrine (Fig. S1H). Similar to the blood pressure results, Y198F arteries phenocopied SMC *Panx1* KO vessels, with decreased phenylephrine-induced contraction compared to controls (Fig. 3B) which remained unchanged with PxIL2P treatment and showed small, but significant differences with SPIR treatment at some doses. S205A arteries showed no differences compared to controls, with phenylephrine contractile responses exhibiting nonsignificant trends to decreases with both PxIL2P and SPIR inhibition (Fig. 3C). Y308F vessels showed the opposite response to Y198F, with slightly enhanced phenylephrine-induced contraction. However, these increases were abolished when Y308F arteries were treated with PxIL2P and SPIR, where large reductions in phenylephrine-induced contraction were observed (Fig. 3D).

**Fig 3.**
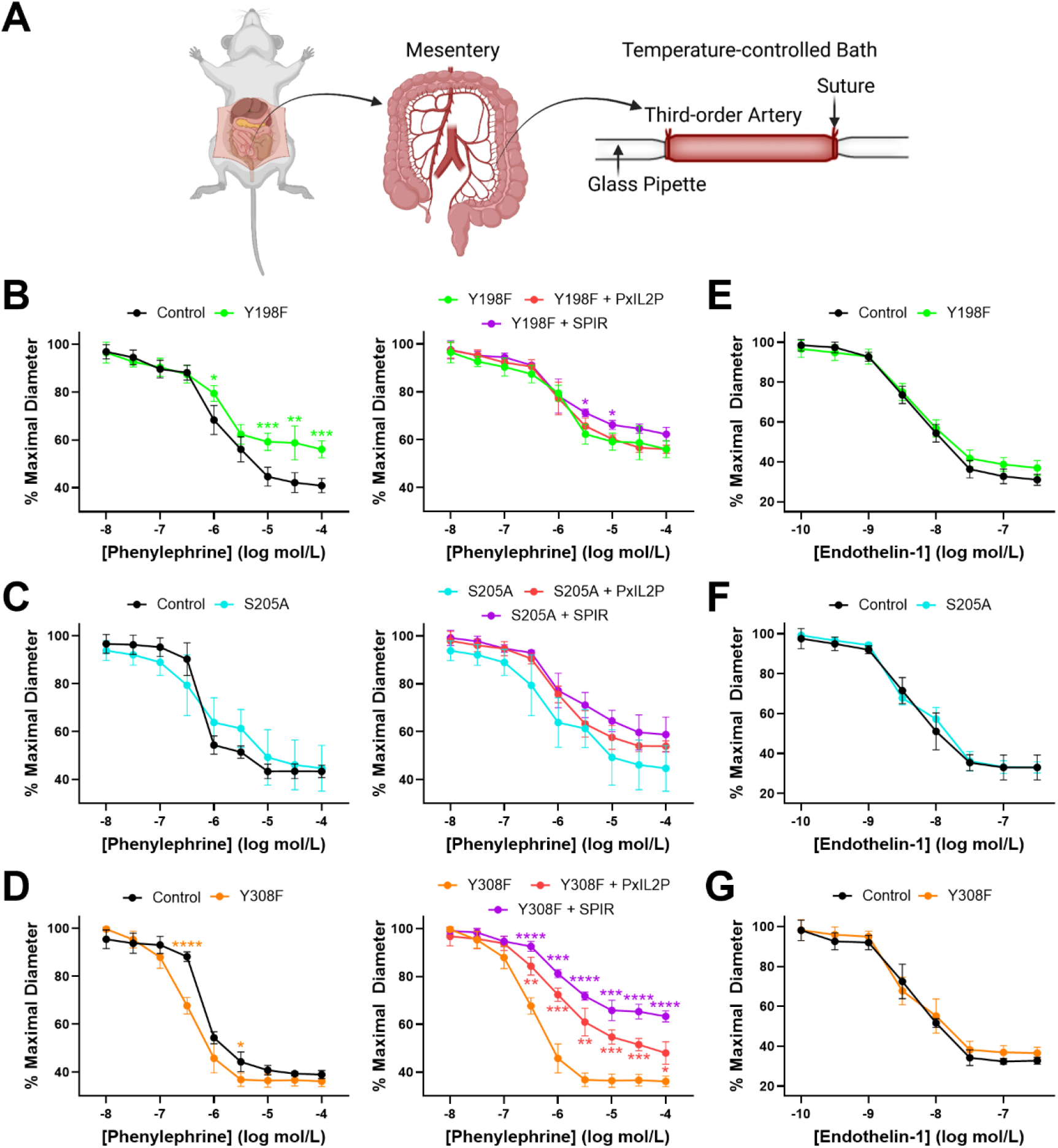
PANX1 Y198F resistance arteries have reduced contractility in response to α-adrenergic stimulation, whereas Y308F vessels exhibit slightly increased responses. (**A**) Schematic for third-order mesenteric artery dissection and pressure myography setup. Created with BioRender.com. Cumulative dose-response curves to increasing concentrations of phenylephrine with and without PANX1 inhibitors PxIL2P or spironolactone (SPIR) (**B**, Y198F; **C**, S205A; **D**, Y308F) or endothelin-1 (**E**, Y198F; **F**, S205A; **G**, Y308F) by pressure myography of third-order mesenteric arteries. Lower % maximal diameter indicates increased contraction. Two-way ANOVA followed by Sidak’s multiple comparisons test for **B-G**. *N*=5 mice. \**P*<0.05, \*\**P*<0.01, \*\*\**P*<0.001, \*\*\*\**P*<0.0001.

The altered vasoreactive responses in SMC-specific *Panx1* KO, Y198F and Y308F mice were found to be specific to α-adrenergic signaling since third-order mesenteric arteries from all genotypes exhibited no changes in endothelin-1-induced vasoconstriction (Fig. S2I-J, Fig. 3E-G). Furthermore, EC and SMC health in all *Panx1* KO and phospho-dead mutant mice remained intact as shown by a lack of differences observed in NS309-induced dilation and KCl-induced constriction, respectively, compared to littermate control vessels (Fig. S2K-L, Fig. S5). These results illustrate that phenylephrine responses are dependent on the phosphorylation site modified on PANX1, but Y198 phosphorylation plays the primary role in regulating α-adrenergic vasoconstriction.

### PANX1 phospho-dead mutant transcriptomics and protein-protein interactors resemble controls

PANX1 influences cellular and organismal phenotypes through its contribution to signaling pathways, interaction with other proteins and/or function as a conduit for small molecules and ions (*45, 46*). To understand the mechanisms responsible for the blood pressure and vessel vasoreactivity phenotypes, we first performed bulk RNA sequencing (RNA-seq) of mesenteric arteries isolated from 8-week-old mice to investigate any differences between the phospho-dead mutant mice and controls at the transcript level (Fig. 4A-C, S6; Data File S1 Tables S1-S4). We found all three phospho-dead mutants had similar transcriptomic profiles compared to controls, with Y198F (Fig. 4A, S6A), S205A (Fig. 4B, S6B) and Y308F (Fig. 4C) mesenteric arteries exhibiting only 3 (2 increased abundance, 1 decreased abundance), 96 (92 increased abundance, 4 decreased abundance) and 0 differentially expressed genes, respectively. *Panx1* mRNA was not significantly different between any phospho-dead mutant and controls in accordance with our RT-qPCR of mesenteric arteries (Fig. 1C). These results resembled bulk RNA-seq comparing WT and global *Panx1* KO mesenteric arteries, which exhibited only 20 differentially expressed genes including the decreased abundance of *Panx1* transcripts in the KO vessels (Fig. S7; Data File S1 Tables S1 and S5). Gene Set Enrichment Analysis of altered S205A pathways (Fig. S8; Data File S1 Table S6) indicated gene sets associated with cell migration, extracellular matrix assembly and organization as well as the regulation of membrane potential were the most positively enriched compared to controls. Due to the low numbers/lack of differentially expressed genes in Y198F and Y308F mesenteric arteries, we were unable to perform pathway analysis for these genotypes. Overall, because only ≤0.005% of transcripts are altered in phospho-dead mutant or *Panx1* KO mesenteric arteries compared to controls, differences at the level of protein interaction and/or channel function are more likely responsible for the vascular phenotypes.

**Fig 4.**
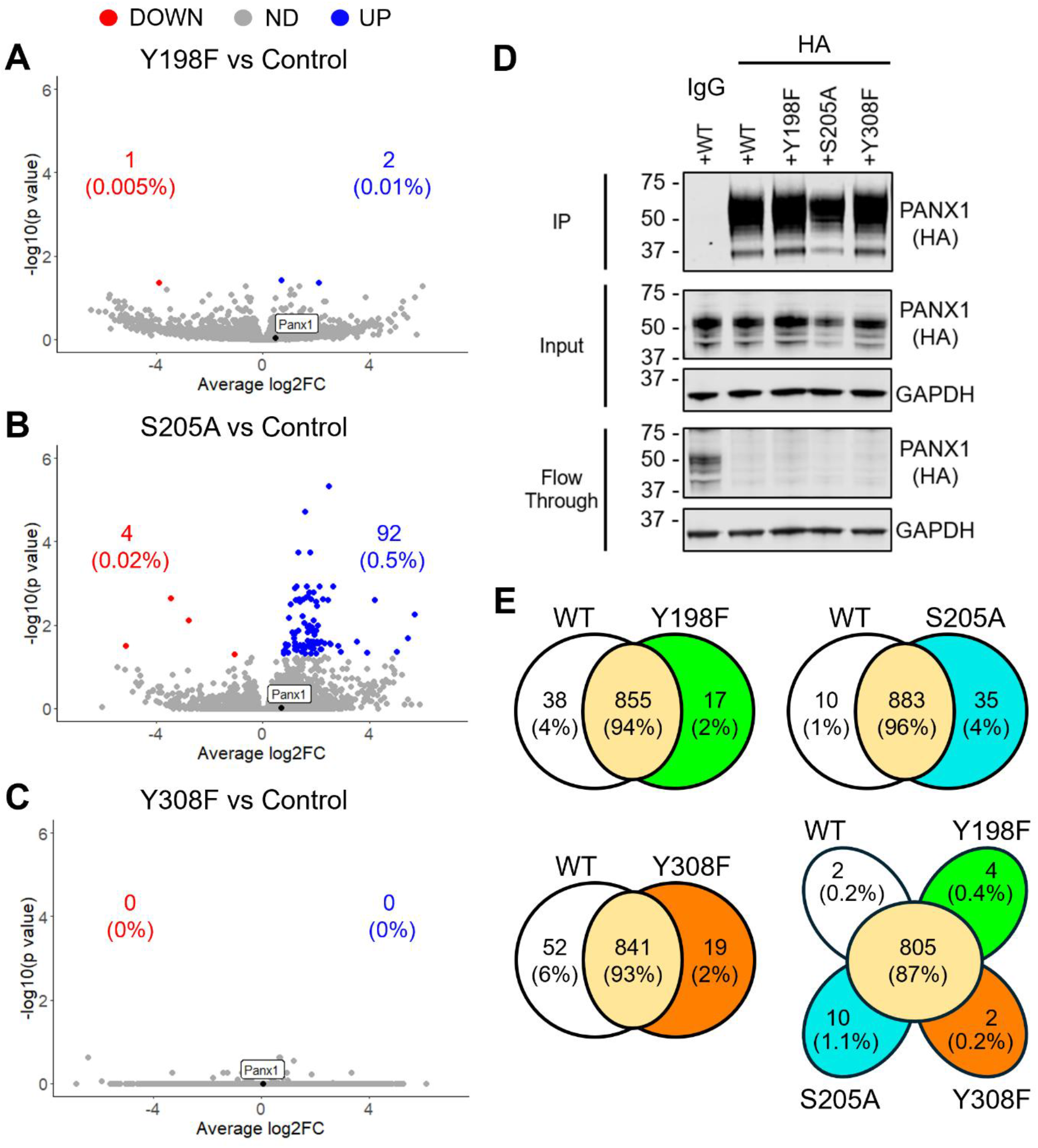
PANX1 Y198F, S205A and Y308F minimally alter transcriptomics in mesenteric arteries or protein interactors in smooth muscle cells. Bulk RNA sequencing of mesenteric arteries from Y198F (**A**), S205A (**B**) and Y308F (**C**) mice compared to controls. Volcano plots exhibit all differentially expressed genes based on *P*<0.05. Blue dots and text show the number and percentage of genes with increased abundance (UP), red dots and text show the number and percentage of genes with decreased abundance (DOWN) and grey dots represent genes not statistically different (ND) from controls. *Panx1* (black dot) levels were unchanged between phospho-dead mutant and control arteries. *N*=3-4 mice. (**D**) *Panx1* knockout (KO) MOVAS were transfected with various PANX1 HA-tagged constructs and immunoprecipitated (IP) using a mouse HA antibody for PANX1 pulldown. Mouse immunoglobulin G (IgG) pulldown of PANX1 wildtype (WT)-transfected *Panx1* KO MOVAS as negative control. GAPDH as a protein loading control for input and flow through, protein sizes in kDa. (**E**) Co-immunoprecipitated lysates from **D** were sent for mass spectrometry to analyze protein-protein interactors of WT and PANX1 phospho-dead mutants. Venn diagrams show the number and percentage of overlapping (yellow) or unique interacting protein clusters for WT (white), Y198F (green), S205A (blue) and Y308F (orange) comparisons. *N*=1 co-immunoprecipitation per construct.

To explore whether PANX1 phosphorylation alters the protein interactome of PANX1, we performed co-immunoprecipitation-mass spectrometry on differentiated *Panx1* KO mouse vascular smooth muscle cells (MOVAS) co-expressing FLAG-tagged α-1DAR and either WT or phospho-dead mutant hemagglutinin (HA)-tagged PANX1 (Fig. 4D-E; Data File S2 Tables S7-9). Successful PANX1 immunoprecipitation for each PANX1 variant was confirmed by Western blot (Fig. 4D), and PANX1 was detected in each sample by mass spectrometry. We also detected known interactors of PANX1 such as actin (*11*) and actin-related protein 3 (*47*) as well as scaffolding proteins AHNAK (*48*) and IQ motif-containing GTPase-activating protein 1 (IQGAP1) (*13*) in all PANX1 co-immunoprecipitations. We identified a total of 1432 proteins in 937 clusters (groups of proteins with highly similar sequences that cannot be uniquely identified based on common peptides present in the analysis). Protein clusters identified for each PANX1 phospho-dead mutant were compared to those of PANX1 WT (Fig. 4E, S9; Data File S2 Tables S10-12). We observed marked overlap between PANX1 Y198F, S205A and Y308F clusters compared to WT (ranging from 93-96%), with a much lower percentage of interactors distinct to each PANX1 variant (ranging from 1-6%). Moreover, when all four PANX1 variant clusters were compared, 87% of clusters were present in all samples and only 0.2-1.1% of proteins were unique to each PANX1 variant, illustrating a high level of similarity between the binding partners of each form of PANX1. EnrichR pathway analysis of protein clusters unique to each PANX1 phospho-dead mutant in comparison to WT showed Y198F, S205A and Y308F did not have any significantly different pathways (Data File S2 Tables S13-18). When WT-specific interactors for each of the PANX1 phospho-dead mutant comparisons were analyzed, WT binding partners compared to both Y198F and S205A were associated with approximately 100 significantly altered pathways. Notable WT versus Y198F pathways were associated with intermediate filament organization, cell fate determination, cell cycle control and muscle contraction. For WT versus S205A, pathways involved in actin assembly, cell cycle regulation and Wnt signaling were observed. However, many of the significant pathways were driven by the presence of 1-2 proteins in the co-immunoprecipitations and were unrelated to SMCs (for example, processes specific to other cell types such as epithelial cells or glia). Lastly, distinct WT versus Y308F interactors were related to telomere maintenance and endocytosis. Thus, since only ≤6% of interactors are altered with PANX1 phospho-dead mutants, and pathways associated with unique interactors are unrelated to SMC contractility or α-adrenergic signaling, differences at the level of protein-protein interactions are not likely responsible for the murine vascular phenotypes.

### Y198F, S205A and Y308F differentially modulate PANX1 channel activity

Structural data has provided evidence that PANX1 channels conduct ions through lateral tunnels, with both ions and metabolites such as ATP moving through their central pore (*8, 10*). Analysis of the Y199, S206, and Y309 human PANX1 residues (corresponding to mouse PANX1 Y198, S205, and Y308, respectively) in the context of the heptamer structure (Protein Data Bank ID: 6WBF (*10*)) revealed the structural and molecular impacts of removing the hydrogen bonding capabilities of each amino acid as well as their ability to be phosphorylated through the missense mutation to either phenylalanine or alanine. Y199 (Fig. 5A), located in intracellular helix 2 (IH2), contributes hydrogen bonding interactions to a cluster of ionic interactions between D138 and K202 on the same subunit and K346’ in the adjacent subunit. These interactions stabilize IH6 and the C-terminal activating domain (CAD, residues 361-370) of PANX1, which is critical for ATP release (*49*). Thus, the Y198F mutation is predicted to impact ATP release. Despite its proximity to Y199 in the peptide sequence, S206 (Fig. 5B) is located away from the pore on the membrane-exposed surface of PANX1. S206 is located in a loop that links IH2 and transmembrane helix S3 and is involved in a network of electrostatic interactions with D138 (IH1), K203 (IH2) and N208 (S3) that positions H134 (a loop between transmembrane helices IH1 and S2). H134, along with R29’ from an adjacent subunit, forms part of the PANX1 ion-conducting side tunnel (*10*). Thus, mutation of this residue is predicted to alter ion conductance. Y309 (Fig. 5C) is located on IH4 and hydrogen bonds to S344, which in turn hydrogen bonds to D147. These interactions stabilize the IH1, IH4, and IH6 bundle that lines the pore (*50*). However, unlike Y199, which interacts with the C-terminal end of IH6, Y309 interacts with the N-terminal end of IH6, which is closer to the ion-conducting side tunnel. Therefore, predicting how a mutation or phosphorylation at Y309 would affect ATP release or ion conductance is difficult. Mutating these phosphorylation sites would prevent the phosphorylation-regulated conformational changes in PANX1, which are likely to regulate the function of the regions of the protein that each site stabilizes.

**Fig 5.**
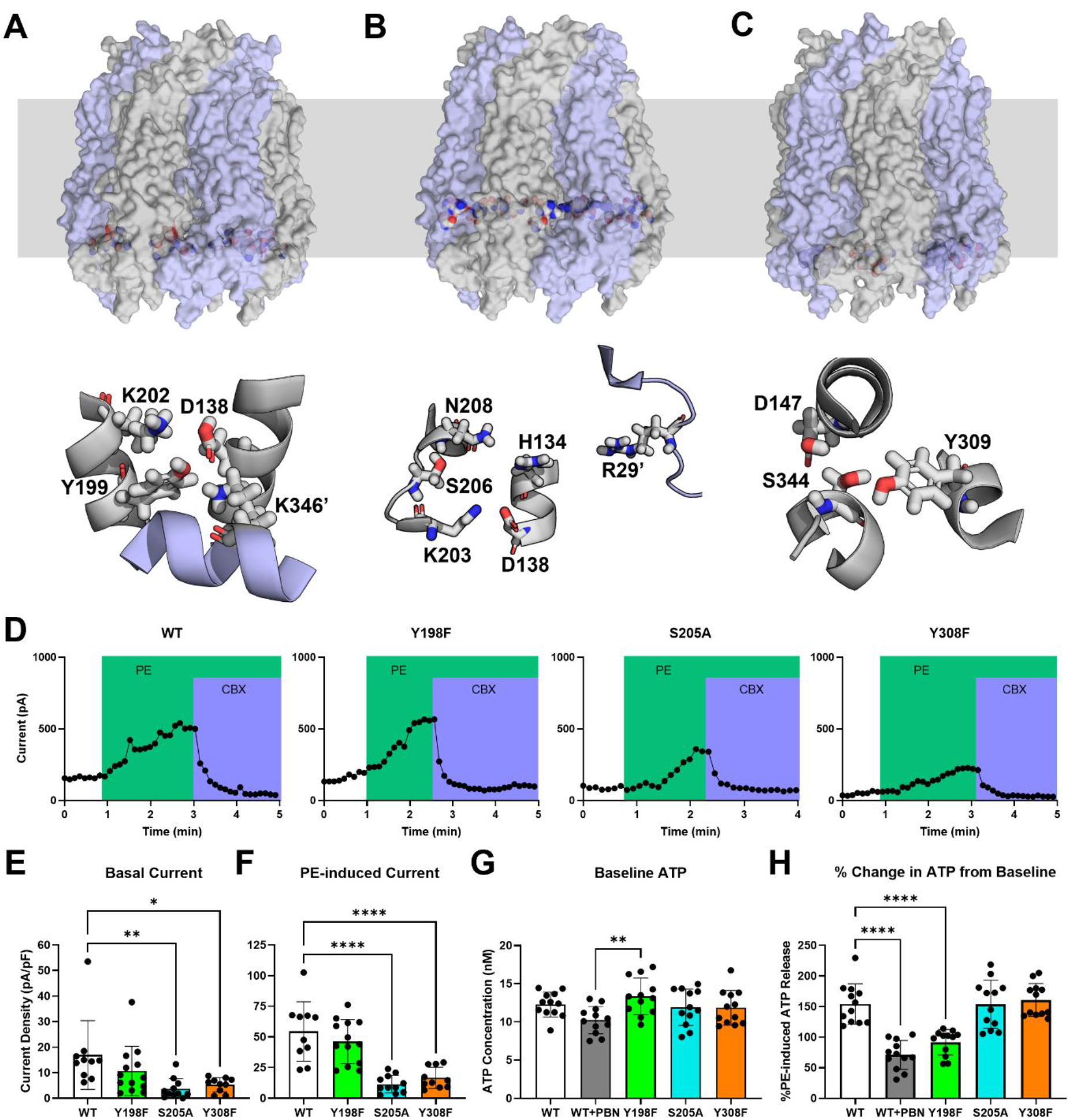
PANX1 Y198F channels have reduced phenylephrine-induced ATP release, whereas S205A and Y308F channels exhibit decreased current. Molecular impacts of human PANX1 structure (Protein Data Bank ID: 6WBF) at sites where mutations Y199F (**A**), S206A (**B**) and Y309F (**C**) were introduced. The context of the mutant and interacting residues (rendered as spheres and colored by element; carbon, gray; nitrogen, blue; oxygen, red; hydrogen, white) within the heptamer is shown with the subunits rendered as surface with alternating light blue and gray coloring. Gray band represents the lipid bilayer. (**A**) Y199 (in intracellular helix 2 (IH2)) hydrogen bonds with D138 and K346’ in the adjacent subunit, stabilizing IH6. (**B**) S206 is involved in a hydrogen bonding network that can position H134 in the putative PANX1 ion-conducting side tunnel (*10*). (**C**) Y309 hydrogen bonds with S344, which interacts with D147, stabilizing the interactions of IH1, IH4, and IH7. Images were made with PyMOL. (**D**) Whole-cell patch-clamp recordings of mouse PANX1 wildtype (WT), Y198F, S205A and Y308F currents over time in human embryonic kidney (HEK293T) cells co-transfected with the mouse α-1D adrenergic receptor and indicated mouse PANX1 construct during phenylephrine (PE, 20 µM) and carbenoxolone (CBX, 20 µM; PANX1 channel blocker) treatment. CBX-sensitive current density of mouse PANX1 WT or phospho-dead mutant channels at baseline (**E**) and after PE stimulation (**F**) (*n*>18 cells, *N*>5 independent transfections for each construct). One-way ordinary ANOVA, followed by Tukey’s multiple comparisons test. Basal ATP concentration (**G**) and %PE-induced ATP release (**H**) (% change from unstimulated/basal conditions to 100 µM PE) from HEK293T cells co-transfected with mouse α-1D adrenergic receptor and indicated mouse PANX1 construct. WT-transfected cells were also inhibited with 1 mM of the PANX1 channel blocker probenecid (WT+PBN). *N*=4 independent experiments, *n*=3 where each dot represents one well. \**P*<0.05, \*\**P*<0.01, \*\*\*\**P*<0.0001. Bars represent mean ± SD.

To corroborate these predicted structural and molecular changes with alterations in channel activity, we assessed both ionic currents and small molecule flux through each specific phospho-dead mutant PANX1 in comparison to WT using HEK293T cells co-transfected with the α-1DAR and each PANX1 variant. All PANX1 phospho-dead mutants showed similar protein expression, glycosylation and trafficking, with PANX1 localizing to the cell surface (Fig. S10). First, we performed whole-cell patch-clamp electrophysiology to analyze basal and phenylephrine-stimulated current in Y198F, S205A and Y308F PANX1 channels (Fig. 5D, Fig. S11). These currents were sensitive to carbenoxolone (CBX) and displayed weakly-rectifying profiles as expected for PANX1. Under unstimulated conditions, basal CBX-sensitive current in Y198F channels resembled WT, whereas basal current was significantly reduced in S205A and Y308F channels (Fig. 5E). Phenylephrine stimulated CBX-sensitive current in all channel constructs (Fig. S11B), but where α-1DAR-activated current was not different between Y198F and WT channels, it was significantly reduced in S205A and Y308F channels (Fig. 5F). Lastly, we used an ATP release assay to investigate the ability of each PANX1 channel to mediate small molecule movement. Basally, ATP release was not different between WT and the individual phospho-dead mutants (Fig. 5G). Likewise, the percent change in ATP release triggered by phenylephrine was not different between WT and either S205A or Y308F channels (Fig. 5H). However, we found that Y198F channels showed significantly lower ATP release with phenylephrine administration, obtaining levels similar to those seen after inhibition of WT channels with the PANX1 inhibitor probenecid (PBN). Altogether, these results illustrate that Y198 phosphorylation regulates PANX1-mediated ATP release, whereas S205 and Y308 phosphorylation influence ion conductance.

## DISCUSSION

Using novel PANX1 phospho-dead mutant mouse models, we uncovered distinct roles of PANX1 Y198, S205 and Y308 phosphosites in the resistance artery vasculature. These roles may be explained by their differential regulation of PANX1 small molecule versus ion channel activity in response to α-adrenergic stimulation as opposed to their effects on mRNA signaling or protein-protein interactions. Ultimately, our findings integrate previous research regarding PANX1 function in the resistance vasculature, phosphorylation and channel activity, providing a multi-scale analysis of how biochemical changes through post-translational modifications can translate into whole body phenotypes.

We illustrated that the PANX1 Y198 phosphosite is a major regulatory feature for α-adrenergic controlled blood pressure and vasoconstriction through stimulation of PANX1-mediated ATP release. Y198F mice phenocopied the reductions in blood pressure and phenylephrine-induced vessel contractile responses seen in SMC-specific *Panx1* KO mice (*16*) and vasoconstriction of Y198F third-order mesenteric arteries showed negligible changes with PANX1 inhibition by spironolactone, further demonstrating the importance of the PANX1 Y198 phosphosite in mediating α-adrenergic effects in the vasculature. Genetic approaches to remove the possibility of PANX1 Y198 phosphorylation in our mice are consistent with what we have seen previously with PxIL2P peptide pharmacological inhibition, which decreased both mean arterial pressure and phenylephrine- or electrical field stimulation-induced vasoconstriction (*15, 16, 19*). We also know through previous work that α-adrenergic vasoconstriction was disrupted by removing extracellular ATP, inhibiting purinergic receptors or blocking PANX1 channel activity (*18, 19*). Considering that ATP release from PANX1 Y198F channels was compromised but remained intact in both S205A and Y308F channels, and Y198 phosphorylation in SMCs was shown to occur through Src family kinases (*17*), it seems likely that PANX1-mediated ATP release is stimulated by Src phosphorylation of PANX1 at the Y198 residue. Thus, Src-mediated Y198 phosphorylation is permissive for α-adrenergic-mediated signaling, triggering the release of ATP through PANX1 channels and downstream purinergic signaling to drive vessel constriction. Furthermore, PANX1 Y198 and Y308 phosphorylation appear to have opposite effects on α-adrenergic vasoconstriction, meaning phosphorylation at the Y308 phosphosite (which has also been shown to occur through Src family kinases (*33, 36*)) may be involved in modulating the channel response to α-adrenergic signaling. The S205/S206 PANX1 phosphosite is not necessary for regulating α-adrenergic-mediated contraction of SMCs, but has more critical roles in other cell types, such as cardiomyocytes and T cells during ischemia-reperfusion and airway epithelium inflammation, respectively (*34, 35*).

Interestingly, both PANX1 S205A and Y308F channels showed deficits in ion conductance but had differing blood pressure and vasoreactivity phenotypes, where there were no differences evident in S205A mice, but Y308F mice exhibited increased MAP and phenylephrine-induced contractile responses. It has been demonstrated that obese humans have increased plasma and urinary norepinephrine levels and heightened sympathetic activity which correlates with visceral adiposity (*51–54*). Consistent with this notion, Y308F mice had significantly higher mass driven by a large increase in percent fat mass, from 13.4% in controls to 21.5% in Y308F mice, as well as a trend to elevated urinary norepinephrine levels compared to WT mice (Fig. S12). PANX1 is present in adipose stromal cells and visceral adipose tissue, where it regulates adipose stromal cell differentiation and fat accumulation, and global *Panx1* KO mice have increased fat mass which was attributed to enhanced adipogenic differentiation of *Panx1* KO adipose-derived stromal cells (*55*). Since the Y308F mutation is present globally, it is possible that the increased blood pressure and contractile responses in Y308F mice may be independent of the resistance vasculature. Subsequent work should investigate the role of Y308 phosphorylation in adipose tissue, as well as obesity development and its relation to hypertension.

Treatment resistant hypertension affects between 100-500 million people worldwide and is associated with increased risk of cardiovascular morbidity and mortality (*56*). When we challenged PANX1 phospho-dead mutant mice with a sympathetic-driven hypertension model, we observed that Y198F mice were protected from blood pressure elevation to the same degree as controls treated with the sympathetic inhibitor clonidine. Surprisingly, Y308F mice did not have elevated blood pressure compared to controls as was observed at baseline, possibly due to a saturating effect of the blood pressure phenotype. Previous work from our group showed hypertensive arteries have markedly increased phosphorylated Y198 PANX1 in SMCs (*17*) and constriction of arterioles isolated from treatment resistant hypertension patients could be reduced by PxIL2P administration (*20*). Collectively, these findings suggest PANX1 Y198 phosphorylation in SMCs is not only a driving factor in blood pressure homeostasis, but its dysregulation may also contribute to the development of hypertension. Moreover, the PANX1 protein sequence shows 94% conservation between humans and mice (*57*), with all three phosphosites conserved (*58, 59*), and overexpression of human PANX1 in the iSMC PANX1 OE mouse model exhibited elevated blood pressure (*19*), exemplifying the potential translatability to our findings. Future studies should evaluate the use of PANX1 Y198 phosphorylation/channel inhibitors as a therapeutic approach alone or in combination with first-line anti-hypertensives, especially in cases of treatment resistant hypertension which remain difficult to manage.

Although individual phosphorylation sites and functional consequences vary by channel family and cellular context, essentially every major class of ion channel has documented regulation by phosphorylation-dependent signaling (*60, 61*). Pannexin channels are no exception, with multiple phosphorylation sites regulating channel activity (*59*). When we analyzed PANX1 phospho-dead mutant channel function, we saw decreased ATP release from Y198F channels compared to WT but conserved channel currents. This matched ATP release and electrophysiology results from phenylephrine-stimulated HEK293 cells treated with PxIL2P (*16*) or expressing PANX1 Y198A (*62*). HEK293 cells expressing the PANX1^YLK>AAA^ mutant plasmid (mouse residues Y198-K200, located on IH2) showed conflicting results, with decreases in both ATP release and current responses. One explanation is that YLK unfolds or partially unfolds IH2, thereby affecting ATP release stabilized by Y198 and the positioning of S205 and other residues important to ion conductance. Contrary to our Y198F results, we observed reduced current in both S205A and Y308F channels after phenylephrine stimulation. This was consistent with reduced current in S205A-expressing HEK293T cells treated with phenylephrine (*34*) and anoxia-induced WT hippocampal pyramidal neurons inhibited with the TAT-Panx_308 p_eptide (*33, 36*). However, our group recently identified ATP release from rat hippocampal neurons, most likely stimulated by non-ionotropic NMDAR signaling and Src-dependent PANX1 Y308 phosphorylation, contributed to late phase long-term depression (*37*). Furthermore, a pre-print investigating Y308 phosphorylation in oocytes using a Y308E phosphomimetic mutation found that Y308E PANX1 channels are constitutively active, conducting both ATP and various ions, but the authors did not investigate Y308F channels (*50*). It is plausible that different cell types and signaling mechanisms could stimulate one type of channel activity over the other, but the structure of PANX1 should be consistent in each condition. However, variations exist in the PANX1 channel cryo-EM structures that have been resolved (*45*) and more work is needed to further understand the intra- and inter-subunit interactions that exist in the PANX1 channel as well as how phosphorylation could affect them.

Our findings also parallel work involving connexin hemichannels in which separate gating mechanisms for small molecule and ion movement were observed (*22*). There have been many instances throughout the pannexin literature in which changes in small molecule movement (as measured by ATP release or dye uptake assays) did not match ion conductance (as measured by electrophysiology) (*63–67*), but we did not have the context for these differences until the presence of side tunnels in the PANX1 channel structure was discovered (*10*). For example, in pulmonary artery SMCs, SPIR and ^10^Panx inhibitory peptide exposure diminished the increase in intracellular calcium normally seen with acute hypoxic stimulus, but ATP release was undetectable. This indicated PANX1 calcium ion conductance rather than ATP release was critical in hypoxic pulmonary vasoconstriction (*68*). Ruan *et al.* provided evidence that ion movement through the alternate side tunnel pathway could occur even when the C-terminal tail blocked the main pore, preventing small molecule movement (*10*). Additionally, it seems that these alternate PANX1 channel compartments may be regulated separately, where the C-terminal tail acts through a ball-and-chain mechanism to plug the central pore while the N-terminus may gate side tunnel ion flux (*10, 24, 25*). Since side tunnels have also been identified in innexins and volume-regulated anion channels, this alternate pathway of ion conductance may be a common structural aspect of large pore channels (*10*). Our study reiterates the independent regulation of each type of PANX1 channel function and points to the contribution of phosphorylation as one mechanism to modulate this activity.

The differential effects of the three phospho-dead mutations on PANX1 channel activity reflect the distinct structural roles of each residue and allow prediction of the roles of phosphorylation. Y199 contributes to stabilizing IH6 and CAD, which is directly implicated in ATP release (*49*). Electrostatically, phosphorylation of Y199 is easily accommodated by the two lysine residues (K202 and K346’) despite the presence of D138. Thus, structural analysis predicts phosphorylation will rearrange the ionic networks to stabilize an open conformation. In contrast, S206 is away from the pore, and both structural and experimental evidence support that phosphorylation at this site regulates the ion-conducting side tunnel. One possible structural rearrangement is that phosphorylation of S206 forms a salt bridge with K203, disrupting the salt bridge with D136 and thereby repositioning IH1. As a result, H134 would move and modulate ion conductance. With respect to Y309, the experimental data indicates that mutation to phenylalanine decreases ion conductance and does not impact ATP release. Combined with the structural analysis, the N-terminal end of IH6 is predicted to be coupled to ion conductance, and the C-terminal end to ATP release. In contrast to the other two sites, there does not appear to be a positive charge near Y309 that would regulate a discrete rearrangement to reposition IH6 or the surrounding helices. Phosphorylation would be repulsive and destabilize the tertiary interactions observed in the structure. Since ATP release is not disrupted by the mutation, IH6 remains well-positioned, whereas interactions with IH5, which forms part of the ion-conducting side tunnel (*10*), may be disrupted. Altogether, these findings suggest that phosphorylation of Y198, S205 and Y308 regulates distinct aspects of PANX1 permeability, with Y198 serving as a selective switch for ATP release and S205 and Y308 contributing to ion conductance.

Despite being canonically known as a channel protein, PANX1 also has channel independent functions in various cell types. PANX1 plays a signaling role in the Hippo (*13*) and Wnt (*12*) pathways, and can interact with many binding partners including actin (*11*), actin-related protein 3 (*47*), calmodulin and β-catenin (*12*) to promote cellular changes such as proliferation and migration. In our study, we observed very few changes in mRNA expression when PANX1 Y198, S205 and Y308 phosphosites were mutated. We would not anticipate considerable transcriptomic changes, since the alteration of only one amino acid to disrupt a phosphorylation site would be more likely to have a greater impact post-translationally on protein activity rather than transcriptional programs. We were surprised that the PANX1 interactome was minimally affected by the inability of PANX1 to be phosphorylated at Y198, S205 and Y308 sites based on previous findings regarding its interaction with caveolin-1 (*15*) and α-1DAR (*18*) in SMCs. Generally, we saw that the core interactome was largely consistent between WT and each phospho-dead mutant PANX1, with the few proteins unique to WT or each of the mutants having little biological relevance to the phenotypes seen in this study. Therefore, changes in PANX1 channel activity were the predominant effect of removing these phosphosites. However, it should be noted that both the transcriptomic and interactome analyses were performed in differentiated but unstimulated conditions, and challenging the vascular SMCs may produce larger variations in binding partners between PANX1 WT and phospho-dead mutants. For example, PANX1 was only seen to interact with caveolin-1 after phenylephrine treatment (*15*). Moreover, some protein associations such as kinase-substrate interactions may be too transient to be detected by co-immunoprecipitation (*69*).

Overall, this work demonstrates the distinct roles for different PANX1 post-translational modifications depending on which physiological process is being regulated, suggesting that interplay between different modifications can enable complementary regulation of PANX1 activity. Future work should assess the physiological consequences of PANX1 phosphorylation sites in other tissue contexts or diseases—such as in ECs where PANX1 regulates leukocyte emigration and venous permeability during inflammatory responses (*29, 40*)—since these phosphorylation sites may have differential effects in other cell types and/or processes regulated by PANX1 currents versus small molecule release.

## MATERIALS AND METHODS

### Ethical approval

All mouse experiments were approved by the University of Virginia’s Animal Care and Use Committee (protocols #3648, #4051), and the Animal Use Subcommittee of the University Council on Animal Care at the University of Western Ontario (protocol #2023-109) and University of Calgary (protocol #AC21-0181). Experiments were conducted in accordance with the National Institutes of Health (NIH) Guide for the Care and Use of Laboratory Animals.

### Mouse models

PANX1 Y198F mutant mice were generated by Ingenious Targeting using a CRIPSR/Cas9 gene editing approach. To target Cas9 to the correct location in the *Panx1* gene and introduce the TAC→TTC point mutation, we used the GGG protospacer adjacent motif (PAM) site, PANXgR2.0R guide RNA 5’-TCAAGTACTGCTCCACGATT-3’ and PANXsD2.0F template DNA 5’-GTCTGTGGGAGATATCTGAAAGCCACTTCAAGTATCCAATTGTCGAACAGTTCTTGAAGACAAAAAAGAACTCTAGTCAT-3’. Mice were genotyped using two polymerase chain reactions (PCRs) to distinguish between WT and Y198F mutant DNA. The mutation-specific PCR used PANX 3.0F 5’-TCAAGTATCCAATTGTCGAACAGTTCT-3’ and PANX 2.0R 5’-TAGCCTTCAGACTTGAAATGTAGAACA-3’ primers, with an expected PCR product size of 402 bp. The WT-specific PCR #1 used PANX 1.1F 5’-CGCAGCAGTCAGATGAAGTTAC-3’ and PANX 2.0R 5’TAGCCTTCAGACTTGAAATGTAGAACA-3’ primers, with an expected PCR product size of 432 bp. F0 founders with mosaicism were bred with WT mice to generate pups with germline transmission. F1 heterozygotes were identified by PCR using the above mutation-specific PCR reaction and the WT #2 reaction with PANX 1.1F 5’-CGCAGCAGTCAGATGAAGTTAC-3’ and PANX 2.0R 5’-TAGCCTTCAGACTTGAAATGTAGAACA-3’ primers (expected PCR product size of 560 bp). For both F0 founders and F1 generation mice, PCR products with mutation-specific PCR products were extracted and sequenced by Inference of CRISPR Edits analysis (Synthego) to confirm the introduced mutation. F1 heterozygotes were then bred with WT C57BL/6N mice purchased from Taconic Biosciences and further propagated by heterozygous crosses.

PANX1 Y308F mice were generated by CRISPR/Cas9 to introduce a TAT→TTT point mutation at base pair 311 in concert with the Clara Christie Centre for Mouse Genomics at the University of Calgary. Two PAM sites, CCG and CCT, were targeted with 5’-GCAGAAAACGGACATTCTCA-3’ and 5’-GCCCACCTTCGATGTTCTAC-3’ guide RNAs which flank the TAT codon, allowing for insertion of the donor template ssODN 5’-TCCAGCTGCTCAGCCTCATTAACCTCATTGTGTATGCTCTGCTGATTCCCGTGGTCGTCTACACGTTCTTCATCCCATTTCGGCAGAAAACGGACATTCTCAAAGTGTTTGAAA TATTGCCCACCTTCGATGTTCTACATTTCAAGTCTGAAGGCTACAATGACTTGAGCCTCTACAACCTTTTTCTGGAAGAGAACATA-3’. Mice were genotyped using PCR with 5’-GTAGAACATCGAAGGTGGGC-3’ and 5’-TGAGAATGTCCGTTTTCTGC-3’ primers and successful point mutation was confirmed by cleavage with SspI to yield two products of 310 bp and 200 bp compared to the single 510 bp product of the WT. Mice were then backcrossed for six generations with WT C57BL/6J mice and heterozygous Y308F breeding pairs were established.

PANX1 S205A mice as well as EC-specific and SMC-specific *Panx1* KO mice were generated previously (*16, 29, 34*). Global *Panx1* KO mice were provided by Genentech, Inc. (*70*). Littermates were used as controls from each corresponding mouse line except global *Panx1* KO mice which were compared to WT C57BL/6N mice purchased from Taconic Biosciences. Mice were housed under a 12-hr light/dark cycle and fed normal chow *ad libitum*. Male mice within the age range of 8-20 weeks were used for all experiments. Genotype percentages from phospho-dead mutant heterozygous crosses only included litters in which all pups reached weaning age. Images of mice were taken on an iPhone 15 (Apple, Inc.).

### Genotyping

Mice were genotyped from tail biopsies using real-time PCR with specific, proprietary probes designed for each mutation (Transnetyx, Inc.). GENEWIZ Sanger sequencing of PCR products generated from tail genomic DNA was used to verify Transnetyx genotyping. Genomic DNA was isolated as described previously (*71*) and PCR products were generated using MyTaq™ Red Mix (Meridian Bioscience, BIO-25043) according to the manufacturer’s protocol for a 20 µL reaction with an annealing temperature of 61°C and 45 sec extension. PCR amplification with primers PMSeq For 5’-CCGCAGCAGTCAGATGAAGT-3’ and PMSeq Rev 5’-AGAGCCCACCCTACCTTTGA-3’ (Integrated DNA Technologies) included *Panx1* exon 4 and the beginning portion of intron 4. To ensure successful amplification, 10 µL of PCR product was separated in a 1% agarose gel containing 1:10,000 SYBR Safe DNA Gel Stain (Invitrogen, S33102). Samples were prepared with 6X Purple Gel Loading Dye (New England BioLabs, B7024S) with Quick-Load 100 bp DNA ladder (New England BioLabs, N3231S). Gels were imaged using the Invitrogen iBright 1500. Sanger sequencing was performed using the forward primer with the remaining 10 µL of PCR product. Sequencing was visualized using SnapGene Viewer 8.2.1 (www.snapgene.com).

### RT-qPCR

Eight-week-old mice were euthanized using 25 µL of 2,2-Dichloro-1,1-difluoroethyl methyl ether (Fisher Scientific, L1728506) in a 15 mL nose cone until toe pinch reflex was lost, and blood was cleared by cardiac perfusion with 20 mL PBS. The mesentery was dissected out and placed in ice-cold Krebs-4-(2-hydroxyethyl)-1-piperazineethanesulfonic acid (Krebs-HEPES; 118.4 mM NaCl, 4.7 mM KCl, 1.2 mM MgSO_4,_ 4 mM NaHCO_3,_ 1.2 mM KH_2P_O_4,_ 10 mM HEPES and 6 mM glucose, 2 mM CaCl_2 a_t pH 7.4) and pinned on Sylgard™ (Electron Microscopy Sciences, 24236-10) plates. Three vascular trees of mesenteric arteries were cleared of surrounding adipose tissue, dissected out and flash frozen in liquid nitrogen. RNA was isolated using the RNeasy MinElute Cleanup Kit (QIAGEN, 74204), followed by cDNA synthesis using SuperScript IV Reverse Transcriptase (Thermo Fisher Scientific Inc., 18090050). TaqMan^®^ Gene Expression Master Mix (Thermo Fisher Scientific Inc., 4369016) and Thermo Fisher Scientific TaqMan^®^ probes *Panx1* (Mm00450900_m1 FAM-MGB, 4331182) and β-2 microglobulin (*B2m*; Mm00437762_m1 VIC PL, 4448485; in-well control) were used for RT-qPCR with 50 ng cDNA per reaction, in a total reaction volume of 20 µL according to the manufacturer’s instructions. Samples were run in triplicate on a CFX Real-Time Detection System (Bio-Rad) for 40 cycles, quantified using the ΔΔCT method (Bio-Rad CFX Manager™ Software) and presented as a fold change relative to the corresponding control average.

### Western blotting

Protein was isolated using 1x radioimmunoprecipitation assay buffer supplemented with P1, P2 and P3 inhibitors (Millipore Sigma, P8340, P5726, P0044), and quantified using a bicinchoninic acid assay (BCA; Thermo Fisher Scientific Inc., 23225). Three vascular trees each of mesenteric arteries were dissected as above from 8-week-old mice and crushed in liquid nitrogen before sonication, or protein was isolated from cultured cells by cell scraping and sonication. Protein (30 µg for mesenteric arteries, 40 µg for transfected cells) was separated on 10% NuPAGE™ Bis-Tris Midi protein gels (Invitrogen, WG1201BOX) for 85 mins at 170 V and transferred for 45 mins at 100 V onto 0.45 µm nitrocellulose membranes (Genesee Scientific, 84-876). Membranes were blocked for 1 h at room temperature with 5% bovine serum albumin (BSA; Fisher Scientific, BP1600-100) tris-buffered saline (TBS), followed by incubation with anti-PANX1 1:1000 (Cell Signaling Technology, 91137, RRID:AB_2800167), rabbit anti-HA 1:1000 (Cell Signaling Technology, 3724S, RRID:AB_1549585) or mouse anti-FLAG 1:1000 (Sigma-Aldrich, F1804, RRID:AB_262044) overnight at 4°C, or anti-glyceraldehyde-3-phosphate dehydrogenase (GAPDH; Thermo Fisher Scientific, MA5-15738, RRID:AB_10977387) at 1:5000 for 1 h at room temperature. IRDye-800CW goat anti-rabbit (LI-COR Biosciences #926-32211, RRID:AB_621843) and IRDye-680RD goat anti-mouse (LI-COR Biosciences, 926-68070, RRID:AB_10956588) secondary antibodies were incubated for 1 h at room temperature. All antibodies were diluted in 5% bovine serum albumin, 0.05% Tween20 TBS. Blots were imaged using the LI-COR Odyssey CLx Infrared Imaging System and quantified in Image Studio™ Software (LI-COR Biosciences).

### En face

*En face* was performed as described previously (*71*). Briefly, the mesentery was dissected out of 13-week-old mice euthanized by CO_2 a_sphyxiation, placed in ice-cold Krebs-HEPES and pinned on Sylgard™ plates. Third-order mesenteric arteries were cleared of surrounding adipose tissue, separated from surrounding arteries, pinned on Sylgard™ squares by 0.013 mm diameter tungsten wire and fixed with 4% paraformaldehyde at 4°C for 15 min. After phosphate-buffered saline (PBS) washes, microscissors (Fine Science Tools, 15000-03) were used to cut arteries longitudinally, and arteries were pinned with tungsten wire such that the endothelial surface was exposed. Arteries were permeabilized in 0.2% Nonidet P-40 PBS at room temperature for 30 min, blocked with 5% BSA in PBS for 1 h, and incubated with anti-PANX1 1:100 (Cell Signaling Technology, 91137) and anti-claudin 5 1:100 (Thermo Fisher Scientific, 35-2500, RRID:AB_2533200) overnight at 4°C. Preparations were washed with PBS, incubated with donkey anti-rabbit Alexa Fluor™ 647 (Thermo Fisher Scientific, A-31573, RRID:AB_2536183) and donkey anti-mouse Alexa Fluor™ 568 (Thermo Fisher Scientific, A10037, RRID:AB_11180865) secondary antibodies (1:400) for 1 h at room temperature, washed in PBS—with the third wash containing 0.1 mg/mL 4′,6-diamidino-2-phenylindole (DAPI; Invitrogen, D1306)—and mounted with ProLong™ Gold Antifade Mountant with DNA Stain DAPI (Fisher Scientific, P36931). Antibodies were diluted in 0.5% BSA PBS. Immunofluorescence images were taken on a FLUOVIEW 3000 (Olympus Life Science) confocal microscope with a 60x oil objective. Images were processed using Fiji (v1.53c) (*72*).

### EchoMRI

Body composition metrics consisting of fat mass, lean mass and water mass were measured for 13-week-old mice by an EchoMRI Body Composition Analyzer (EchoMRI™). Total body mass was determined using a scale (Fisher Science Education, Model CLF 201). Percent fat, lean and water mass were calculated by normalizing to total body mass.

### Radiotelemetry

Blood pressure was measured by radiotelemetry (Data Sciences International, HD-X10 or PA-C10) in 10-week-old mice, where radiotelemeter catheters were implanted into the left carotid artery such that the probe reached the aortic arch and the battery pack for each probe was secured in a subcutaneous pocket on the right side of the mouse flank. Throughout the surgical procedure mice were fully anesthetized with 1.7-2% isoflurane (Covetrus, 11695067772), and for pain management, buprenorphine HCl (Covetrus, 42023017905) was used as an analgesic before surgery at 0.012 mg/mouse or after surgery as needed. Mice recovered for a minimum of 5 days before baseline blood pressure (systolic, diastolic and mean arterial pressure) and heart rate was recorded in live, freely moving mice for five continuous days using Dataquest A.R.T 20 software (Data Sciences International). Parameters were sampled every minute throughout the recording period, and measurements within each time frame (day 7:00 a.m. to 6:59 p.m, night 7:00 p.m. to 6:59 a.m.) over the five-day period were averaged per mouse.

To induce sympathetic-driven hypertension, phospho-dead mutant and control mice were placed on a 6% HSD (ssniff Spezialdiäten GmbH, S3544-E046) at 8 weeks of age, radiotelemeters were inserted as described above at 10 weeks and AngII-loaded (Sigma, A9525) osmotic pumps (Alzet, 1004) were inserted at 14 weeks. AngII was infused at a rate of 1.0mg/kg/day for 2 weeks, and blood pressure was monitored 5 days after osmotic pump insertion until endpoint at 16 weeks. For clonidine treatment, HSD/AngII mice received either 0.05 mg/kg clonidine hydrochloride dissolved in 0.9% saline or 0.9% saline vehicle control by intraperitoneal injection at 16 weeks. Blood pressure was measured 90 minutes after injections during daytime hours based on previous experimental design (*20*).

### Pressure myography

Individual, intact third-order mesenteric arteries (dissected as described above from 12-week-old mice) were cannulated on glass pipette tips using super fine forceps (Fine Science Tools, 11254-20) and nylon sutures (Danish Myo Technology, P/N 100115) in an arteriograph chamber (Danish Myo Technology) containing cold Krebs-HEPES. To clear any remaining blood, Krebs-HEPES buffer was gently pushed through the artery before cannulating the second side. Arteries were equilibrated over a 30 min period by increasing the pressure from 20 to 80 mmHg in 20 mmHg increments (Big Ben Sphygmomanometer), and heating to 37°C. Cumulative concentrations of endothelin-1 (Sigma-Aldrich, E7764) or phenylephrine (Sigma-Aldrich, P-6126), with and without PANX1 inhibitors PxIL2P (5 μmol/L) (*16*) or SPIR (80 μmol/L) (*20*), were applied to the vessels. Buffer was circulated between the arteriography chamber and a beaker containing an excess reservoir of buffer by using a peristaltic pump (Atalyst Masterflex, FH30) to prevent overheating and facilitate the delivery of pharmacological agents. The Danish Myo Technology cell culture pressure arteriography setup and MyoView software (2015 release) were used to measure vessel diameter (including the lumen, excluding the vessel wall). Following the phenylephrine dose responses, SMC and EC function was evaluated using 30 mM KCl (Fisher Scientific, P217) and 1 μM NS309 (MilliporeSigma, N8161), respectively. Maximal diameter measurements were determined after treating arteries with calcium-free buffer following experimentation and stabilization at a pressure of 80 mmHg. Representative traces for Y198F, S205A and Y308F are found in Fig S4.

### Bulk RNA sequencing

Bulk RNA-seq (Novogene Corporation Inc.) was performed on RNA isolated from Y198F, S205A, Y308F and global *Panx1* KO mesenteric arteries (three vascular trees each) and corresponding littermate or WT controls. Mesenteric arteries were dissected and RNA was isolated as described above from mice at 8 weeks of age. Paired-end 150 base pair reads were generated using the Illumina NovoSeq platform (Novogene Corporation Inc.) and mapped to the Genome Reference Consortium Mouse Build 39 reference genome (GRCm39; Accession ID GCA_000001635.9) using STAR software (v2.7.9a) (*73*). In RStudio (v4.4.2) (*74*), gene counts were generated using featureCounts (*75*) in the Rsubread (v2.20.0) software package (*76, 77*). Differentially expressed genes (DEGs, *P*adj < 0.05) were identified using DESeq2 (v1.46.0) (*78*), and the org.Mm.eg.db package was used to convert Ensemble IDs to gene symbols. Gene Set Enrichment Analysis (*79*) was used to investigate pathway changes. All RNA-seq data can be found in Data File S1. Sequencing data was deposited in the National Center for Biotechnology Information’s Gene Expression Omnibus (GEO; accession number GSE335251).

### Cell culture

*Panx1* KO MOVAS (*80*) were generated previously by targeting exon 2 with CRISPR/Cas9 (WT MOVAS were a kind gift from Dr. Lian-Wang Guo at the University of Virginia) and cultured in 1x Dulbecco’s Modified Eagle Medium (DMEM), high glucose, GlutaMAX™ Supplement, pyruvate (Gibco, 11965092) supplemented with 10% fetal bovine serum (FBS; R & D Systems, S12450) and 0.2 mg/mL Geneticin™ Selective Antibiotic (G418 Sulfate; Fisher Scientific, 10131035), and differentiated for 24 h in 1x DMEM GlutaMAX™ media with 0.25% FBS. HEK293T cells were cultured in 1x DMEM, high glucose media (Gibco, 11965092) supplemented with 10% FBS and 1% penicillin-streptomycin (Gibco, 15140122). Cells were passaged using 1x Trypsin-ethylenediaminetetraacetic acid (0.5%), no phenol red (10x stock diluted with 1x PBS; Gibco, 15400054) after one PBS wash.

### Plasmids

Mouse HA-tagged PANX1 WT, Y198F and FLAG-tagged α-1DAR plasmids were generated previously (*16*). PANX1 S205A and Y308F HA-tagged plasmids were generated by GenScript site-directed mutagenesis from the WT construct. Plasmids were verified by Plasmidsaurus sequencing.

### Co-immunoprecipitation-mass spectrometry

Two million MOVAS cells were seeded in 10 cm plates, with two plates per plasmid. Twenty-four hours after seeding, the media was replaced with 0.25% FBS differentiation media after one PBS wash and cells were transfected with 5.5 µg each of α-1DAR and PANX1 WT or phospho-dead mutant plasmid using Lipofectamine 3000™ (Thermo Fisher Scientific, L3000015) per manufacturer’s protocol 1 h later. Transfected cells were incubated in 0.25% FBS media for 24 h. Protein was isolated from each plate using Pierce™ IP Lysis Buffer (Thermo Fisher Scientific, 87787) supplemented with P1, P2 and P3 inhibitors and cell scraping and quantified by BCA as above. Each plate was used to make 1 mg aliquots for immunoprecipitation and 40 µg aliquots for the input Western blot.

Pierce™ Protein G magnetic beads (Thermo Fisher Scientific, 88847) and the DynaMag™-2 magnet were used for co-immunoprecipitations following the manufacturer’s protocol, except only 25 µL of beads were used per sample and the lysate and the antibody-bound beads were incubated for 30 min at room temperature. Co-immunoprecipitations for each PANX1 variant were run in duplicate using 5 µg of mouse anti-HA (Invitrogen, 26183, RRID:AB_2533049) per immunoprecipitation, with one co-immunoprecipitation taken for mass spectrometry and the other used to confirm the successful capture of the PANX1 bait protein by Western blot. As a pull-down control, 5 µg of mouse immunoglobulin G (IgG) Isotype Control antibody (Thermo Fisher Scientific, 02-6502) was incubated with PANX1 WT lysates. For the Western blot samples, following lysate incubation with the antibody-bound beads, the supernatant was collected to use for flow through. Antibodies and bound proteins were dissociated from beads by incubation with 40 µL of 2x Laemmli sample buffer (Bio-Rad Laboratories, 1610737) supplemented with 10% β-mercaptoethanol for 10 min at 70°C. The input lysate, immunoprecipitation and flow through were immunoblotted using the rabbit anti-HA and anti-GAPDH antibodies as described above.

Mass spectrometry was performed with on-bead digestion at the University of Virginia’s Biomolecular Analysis Facility. Samples were processed following BAF_Protocol_002 On-Bead Digestion (Magnetic Beads) of Proteins in Solution or Provided Already on Beads steps 13-18 (*81*) using 0.1 µg of trypsin for digestion. Peptides were extracted following BAF_Protocol_001 In-gel Digestion steps 19-24 (*82*), desalted and purified as described in BAF_Protocol_003 Desalting with C18 ZipTips (*83*). Liquid chromatography-tandem mass spectrometry was performed on the peptides using an Orbitrap Exploris 480 Mass Spectrometer (Thermo Fisher Scientific, BRE725533) with injection of 7/20 µL following BAF_Protocol_004 LC-MS(/MS) nLC EASY LC1200 and Orbitrap Exploris 480 (*84*). Raw files were submitted to the Mouse_Uniprot_20250613.fasta database search using Proteome Discoverer software (v2.5.0.400) and the results were loaded into Scaffold Q+S software (v5.3.4) according to BAF_Protocol_005 Database Search Proteome Discoverer into Scaffold (*85*) using - 10 ppm precursor, 0.02 Da fragments, carbamidomethyl cysteine fixed, oxidized methionine variable and two unique peptides. Identified protein interactors were exported from Scaffold Q+S software (Data File S2 Tables S7-12) to identify overlap between PANX1 WT and phospho-dead mutant datasets using the dplyr (v1.1.4) (*86*) and tidyverse (v2.0.0) (*87*) packages in RStudio. Pathway analysis on unique interactors in WT versus each phospho-dead mutant comparison (Data File S2 Tables S13-18) was completed using EnrichR (v3.4) (*88–90*), where pathways were considered significant based on an adjusted *P* value < 0.05. Transcription, translation and RNA splicing, processing and stability related pathways were excluded from analyses since these results were most likely a byproduct of PANX1 transient overexpression in MOVAS. Percentages of unique or common interactors were determined by dividing each value by the total number of unique proteins identified in the genotypes being compared. The mass spectrometry proteomics data have been deposited to the ProteomeXchange Consortium using the Proteomics Identifications Database (PRIDE) partner repository (*91, 92*) with the dataset identifier PXD080001 and 10.6019/PXD080001.

### Human PANX1 in silico visualization

Human PANX1 (Protein Data Bank ID 6WBF (*10*)) was rendered to highlight the region of the structure impacted by each phospho-dead mutant with Pymol (The PyMOL Molecular Graphics System, Version 3.0 Schrödinger, LLC.).

### Immunocytochemistry

Coverslips were coated with 0.01% poly-L-lysine (Fisher Scientific, A005C) for 1 h at 37°C in a 6-well plate before 400,000 HEK293T cells were seeded. The media was changed to 1x DMEM media without supplements after one 1x PBS wash 23 h after seeding, and cells were transfected 1 h later with 2 µg each of α-1DAR and the corresponding PANX1 WT or phospho-dead mutant construct following Lipofectamine 3000™ instructions. Four h after transfection, the media was replaced with 1x DMEM supplemented with 10% FBS and 1% penicillin-streptomycin. Coverslips were fixed with ice-cold methanol:acetone (80:20, vol/vol) for 15 min at 4°C 24 h after transfection, and then moved to a humidity chamber where they were blocked with 10% goat serum (Jackson Immunoresearch Inc., 005-000-121) in 1x Dulbecco’s phosphate-buffered saline (DPBS) without Ca^2+^ and Mg^2+^ (Gibco, 14190136) for 1 h at room temperature. Coverslips were incubated with rabbit anti-HA 1:500 and mouse anti-FLAG 1:100 primary antibodies overnight at 4°C. After 1x DPBS without Ca^2+^ and Mg^2+^ washes, coverslips were incubated with Alexa Fluor™ 647 goat anti-rabbit (Invitrogen, A-21246, RRID:AB_2535814) and Alexa Fluor™ 488 goat anti-mouse (Invitrogen, A-11001, RRID:AB_2534069) secondary antibodies for 1 h at room temperature. All antibodies were dissolved in 1% goat serum in 1x DPBS without Ca^2+^ and Mg^2+^. Following 1x DPBS without Ca^2+^ and Mg^2+^ washes, coverslips were stained with 1:500 DAPI (Invitrogen, D1306) in 1x DPBS without Ca^2+^ and Mg^2+^ for 5 min at room temperature and then washed again with deionized water before mounting with ProLong™ Gold Antifade Mountant with DAPI (Thermo Fisher Scientific, P36931). Coverslips were imaged on an Olympus FV3000 as Z stacks with the 60x oil objective and images were processed in Fiji (*72*).

### ATP release

ATP release assays using HEK293T cells were performed based on a combined protocol from previous studies (*29, 93*). Twenty-four-well plates were coated with 0.01% poly-L-lysine for 1 h at 37°C prior to seeding 100,000 HEK233T cells per well. After one PBS wash 23 h after seeding, cells were serum-starved for 1 h, and then transfected with 0.4 µg of α-1DAR and 0.4 µg of PANX1 WT or phospho-dead mutant plasmid (in triplicate) per well using the Lipofectamine 3000™ manufacturer’s protocol. The media was switched back to complete DMEM media 4 h after transfection and ATP was measured 24 h after transfection. Cells were washed once with serum- and phenol-free 1x DMEM (Gibco, 31053028) to remove any residual FBS before incubating for 30 min at 37°C and 5% CO_2 w_ith serum and phenol-free DMEM supplemented with 1% BSA (filter sterilized) to allow for the degradation of any ATP released during the washes. Endogenous ecto-nucleotidases were inhibited by incubating cells with 300 mM ARL 67156 (Fisher Scientific, 12-831-0) for 30 min at 37°C and 5% CO_2,_ and media was collected and transferred to pre-chilled 1.7 mL tubes for basal ATP release. Cells were then stimulated with 100 µM phenylephrine and 300 mM ARL 67156 for 5 min at 37°C and 5% CO_2._ Media was collected and transferred to pre-chilled 1.7 mL tubes to measure phenylephrine-induced ATP release. All treatments were prepared in serum- and phenol-free DMEM and throughout the assay culture media was removed gently from each well using a P1000 pipette. Basal and phenylephrine-induced media was centrifuged at 800 g for 5 min at 4°C to remove any cell debris, and ATP concentrations were determined using the ATP Determination Kit (Thermo Fisher Scientific, A22066) and white 96-well plates (Corning Incorporated, 3912) following the manufacturer’s protocol for 100 µL reactions. Luminescence was measured on a FLUOstar Omega plate reader (Gain 3600, lens emission filter, 0.5 s measurement interval time) 5 min after adding the reaction solution with a multichannel pipette in order to capture the peak of the luciferin/luciferase reaction. After subtracting the blank value from all ATP measurements, the concentration of ATP was determined based on the standard curve and the percent change in ATP from baseline was calculated by the following equation: [(ATP_PE-_ATP_B_)/ATP_B]_*100 (where ATP_PE i_s the phenylephrine-induced ATP concentration, ATP_B i_s the basal ATP concentration).

### Patch-clamp Electrophysiology

HEK293T cells (passages 13–17) were seeded one day prior to transfection at a density of 0.5×10^6^ cells per 35 mm dish in 2 mL of complete DMEM, achieving 70–80% confluency at the time of transfection. Plasmid DNA (3.5 μg each of α-1DAR and PANX1 WT or phospho-dead mutant) was diluted in 200 μL of Opti-MEM, and polyethylenimine (PEI; Polyscience, 26008) was added at a 2:1 (w/w) PEI:DNA ratio (7.0 μg PEI). The mixture was gently pipetted and incubated at room temperature for 15 min to allow complex formation. PEI–DNA complexes were added dropwise to the cells with gentle mixing, and the cells were returned to the incubator (37°C, 5% CO_2_) for overnight culture. For electrophysiological recordings, cells were trypsinized 15 h post-transfection, replated onto glass coverslips and allowed to adhere for 3 h before transferring the coverslips to a recording chamber for patch-clamp experiments. Cells were incubated with a caspase inhibitor 20 μM Q-VD-OPh hydrate (Millipore Sigma, SML0063) and a PANX1 channel blocker 25 μM Trovan mesylate (MedChem Express, HY-103399) during and after transfection in all experiments to prevent inadvertent C-terminal cleavage of PANX1 by endogenously activated caspases and block any constitutive activity from the transfected PANX1 channels, respectively.

Whole-cell voltage-clamp recordings were performed at room temperature using an Axopatch 200B amplifier (Molecular Devices), controlled by pCLAMP 9 software and digitized with a Digidata 1322A (Molecular Devices). Recording pipettes (3–5 MΩ) were fabricated from borosilicate glass capillaries (World Precision Instruments) using a P-97 Flaming/Brown micropipette puller (Sutter Instrument) and fire-polished before use. Cells were bathed in an external bath solution containing 140 mM NaCl, 3 mM KCl, 2 mM MgCl_2,_ 2 mM CaCl_2,_ 10 mM HEPES and 10 mM glucose (pH 7.3; ∼300 mOsm). After formation of high-resistance seals (≥10 GΩ) and establishment of the whole-cell configuration, cells were perfused with bath solution containing phenylephrine (20 µM). Following stabilization of channel activity at +80 mV, the solution was exchanged for bath solution containing 20 µM phenylephrine and 100 µM CBX (Sigma-Aldrich C4790). The internal (pipette) solution contained 100 mM CsMeSO_4,_ 30 mM TEACl, 4 mM NaCl, 2 mM MgCl_2,_ 0.5 mM CaCl_2,_ 10 mM HEPES, 10 mM EGTA, 3 mM Mg-ATP and 0.3 mM Tris-GTP (pH 7.3; ∼290 mOsm). Voltage ramp protocols (−130 to +80 mV; 0.2 mV/ms) were applied every 7 s. CBX-sensitive current was defined as the difference in current at +80 mV before and after CBX application and phenylephrine-induced current was quantified as the change in CBX-sensitive current at +80 mV before and after phenylephrine application. Currents were normalized to cell capacitance to obtain current density.

### Urinary norepinephrine measurements

Norepinephrine was measured in the urine of WT and phospho-dead mutant mice as done previously (*19*). Briefly, after a two-week adjustment period, urine was collected from mice in metabolic cages, and the concentration of norepinephrine was determined by enzyme-linked immunosorbent assay (Novus Biologicals, KA1877).

### Statistics

Data was analysed using GraphPad Prism version 10 with tests specified in figure legends, except when comparing genotypes from heterozygous crosses where Graphpad’s online Chi-square test (https://www.graphpad.com/quickcalcs/chisquared1/) was used.

## SUPPLEMENTARY MATERIALS

Figs. S1-S12

Data files S1, S2

## Acknowledgements

We would like to thank the University of Virginia’s Center for Comparative Medicine for their care of the mice (especially Lauren Lassere), and Dr. Nicholas Sherman (University of Virginia) for his help with the mass spectrometry analysis. We thank Anita Impagliazzo Medical Illustration for creating the graphics for Fig. 1A.

## Author contributions

B.L.O’D., L.S.D., M.B., M.K., and B.E.I. conceptualized the study, B.L.O’D., L.S.D., X.Z., S.A. Loeb, Z.J.J., W.J.S., S.A. Leonhardt., A.G.W., M.A.L., S.C.B. and M.D.W. performed experiments and analyzed data. A.K.J.B., A.K.B., S.R.J., S.P., A.F.T., R.J.T, and D.A.B. provided resources and/or scientific expertise. B.L.O’D., S.C.B., L.C., D.A.B. and B.E.I. wrote the manuscript and B.L.O’D., L.S.D., S.C.B. and L.C. designed figures. All authors reviewed the manuscript.

## Data and materials availability

RNA-seq and mass spectrometry files were uploaded to GEO (accession #GSE335251) and PRIDE databases (accession #PXD080001), respectively, for public availability. All other data needed to evaluate the conclusions in the paper are present in the paper or the Supplementary Materials.

