## Supplemental Figures for "Pannexin 1 phosphorylation sites differentially modulate channel activity and physiological outcomes"

### SUPPLEMENTARY MATERIALS

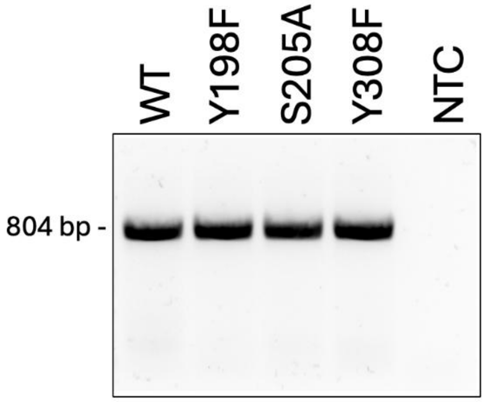


**Fig S1. *Panx1* PCR products sent for Sanger sequencing.** Representative agarose gel of PCR products amplified from wildtype (WT), Y198F, S205A and Y308F genomic DNA isolated from tail snips. All lanes except for non-template control (NTC) show one expected band at 804 bp. *N*=1 mouse.

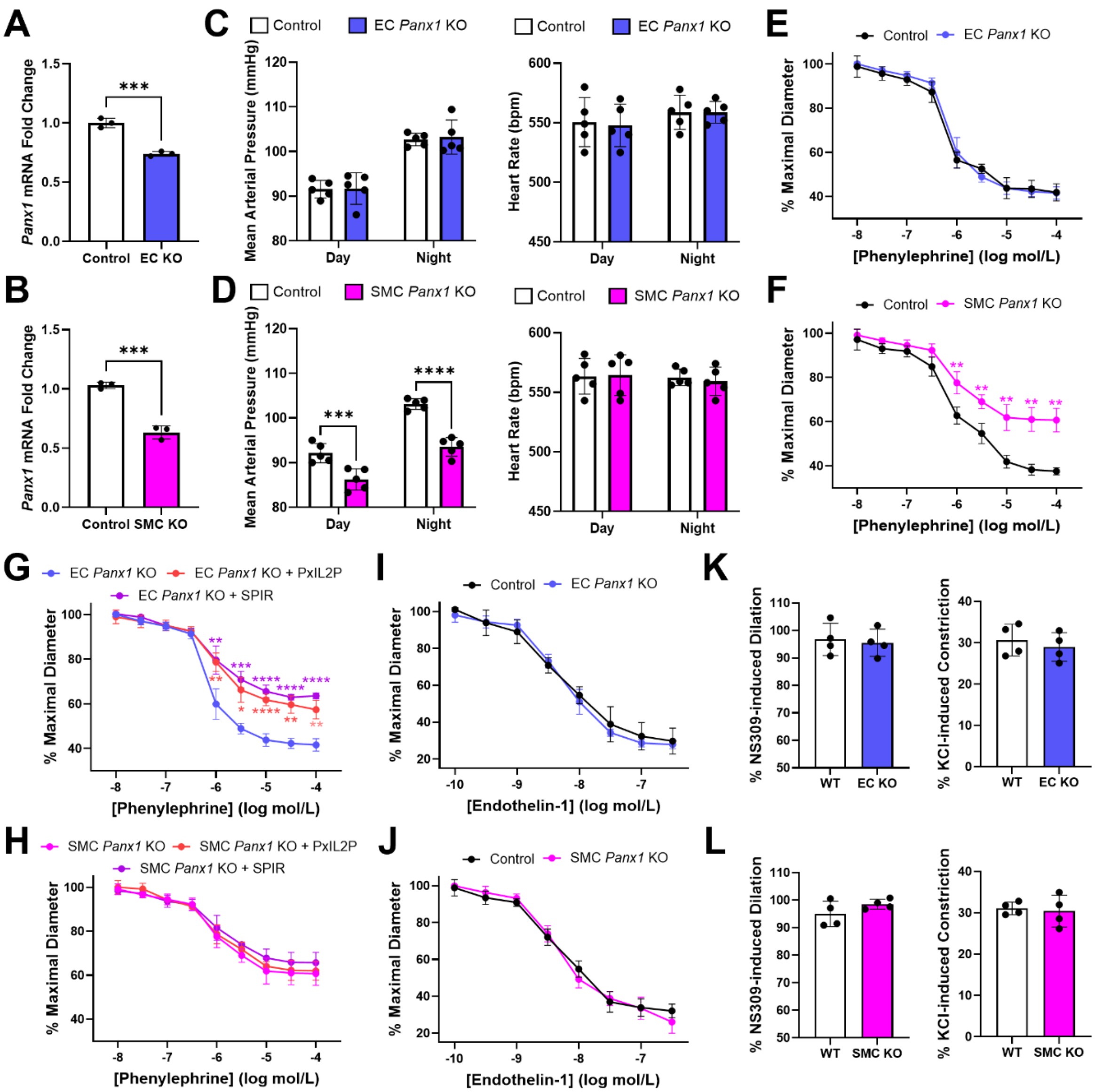

**Fig S2. SMC-specific *Panx1* KO mice have reduced blood pressure and phenylephrine-induced vasoconstriction, which is not seen with EC-specific *Panx1* KO.** RT-qPCR of *Panx1* levels in mesenteric arteries from endothelial-specific (EC KO; **A**) or smooth muscle cell-specific (SMC KO; **B**) *Panx1* knockout and littermate controls. Unpaired t-test. *N*=3 mice. Radiotelemetry-measured baseline mean arterial blood pressure and heart rate during the day and night for EC KO (**C**) and SMC KO (**D**) mice compared to controls. *N*=5 mice. Unpaired t-test within each time point. Cumulative dose-response curves to increasing concentrations of phenylephrine (**E**, EC KO; **F**, SMC KO), with PANX1 inhibitors PxIL2P or spironolactone (SPIR) (**G**, EC KO; **H**, SMC KO) or endothelin-1 (**I**, EC KO; **J**, SMC KO) via pressure myography of third-order mesenteric arteries. Lower % maximal diameter indicates increased contraction. Two-way ANOVA followed by Sidak's multiple comparisons test for **E-J**. Percent dilation to 1 μM NS309 and % constriction to 30 mM KCl for EC KO (**K**) and SMC KO (**L**) third-order mesenteric arteries. Unpaired t-test. **$P$<0.01, ***$P$<0.001, ****$P$<0.0001. Bars represent mean ± SD. *N*=5 mice.

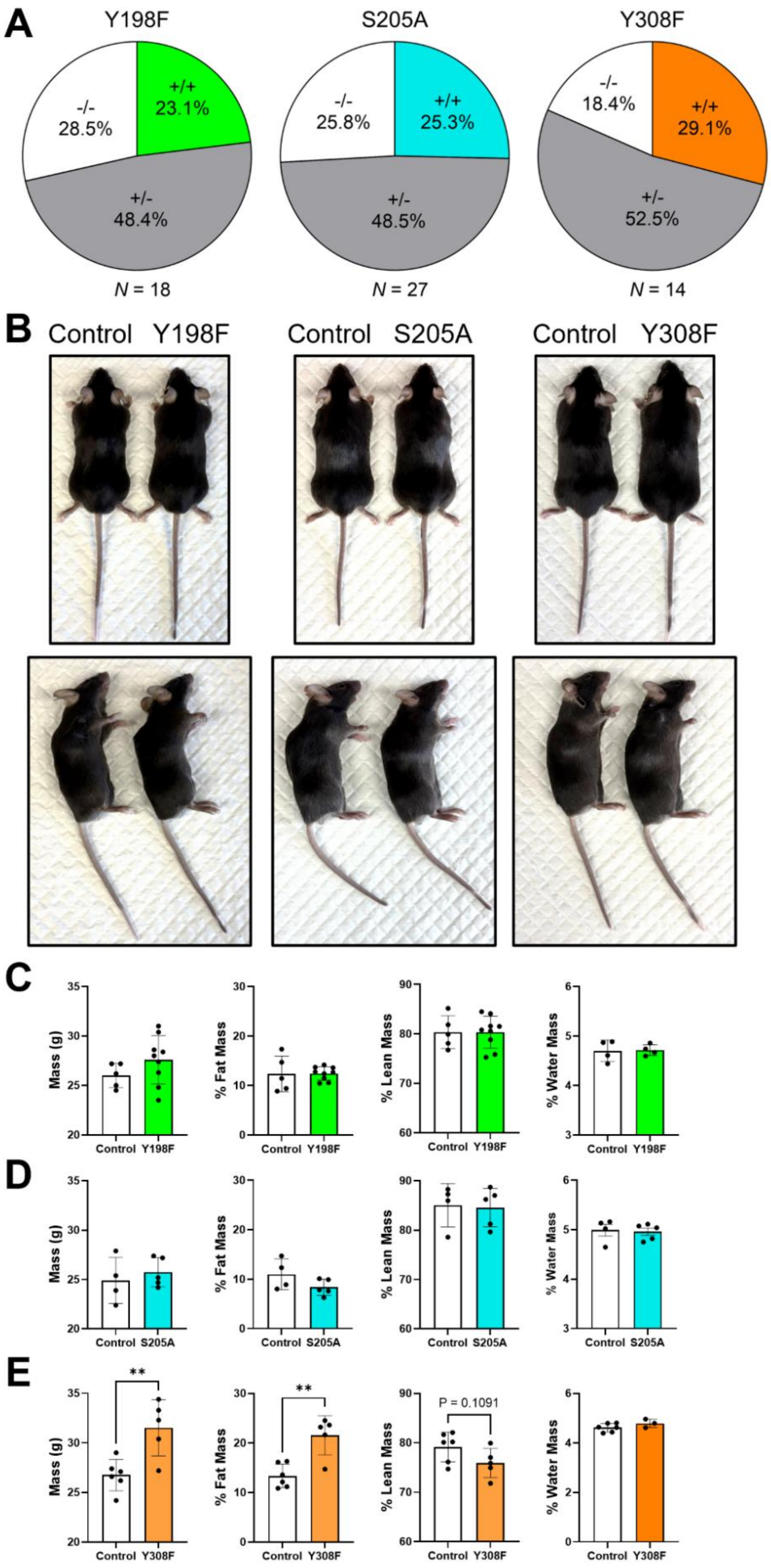
A
Y198F
-/-
28.5%
+/+
23.1%
+/-
48.4%
N = 18
S205A
-/-
25.8%
+/+
25.3%
+/-
48.5%
N = 27
Y308F
-/-
18.4%
+/+
29.1%
+/-
52.5%
N = 14
B
Control Y198F
Control S205A
Control Y308F
C
Mass (g)
% Fat Mass
% Lean Mass
% Water Mass
Control Y198F
D
Control S205A
E
**
P = 0.1091
Control Y308F

**Fig S3. Characterization of PANX1 Y198F, S205A and Y308F phospho-dead mutant mice.** (**A**) Average percentage of homozygous control (-/-), heterozygous (+/-) or homozygous phospho-dead mutant (+/+) pups born per litter from heterozygous crosses. Genotypes were not significantly different from expected Mendelian ratios based on Chi-square goodness of fit test (Y198F $P$=0.7631, S205A $P$=0.8986, Y308F $P$=0.3132). $N$ values of litters analyzed noted in the figure. (**B**) Representative images of control and Y198F, S205A or Y308F male mice at 13 weeks of age. EchoMRI analysis of total body mass, % fat mass, % lean mass and % water mass of 13-week-old male Y198F (**C**), S205A (**D**) and Y308F (**E**) mice compared to controls. Percent masses normalized to total body mass. Unpaired t-test. $^{**}P<0.01$, $N$=3-9 mice. Bars represent mean ± SD.

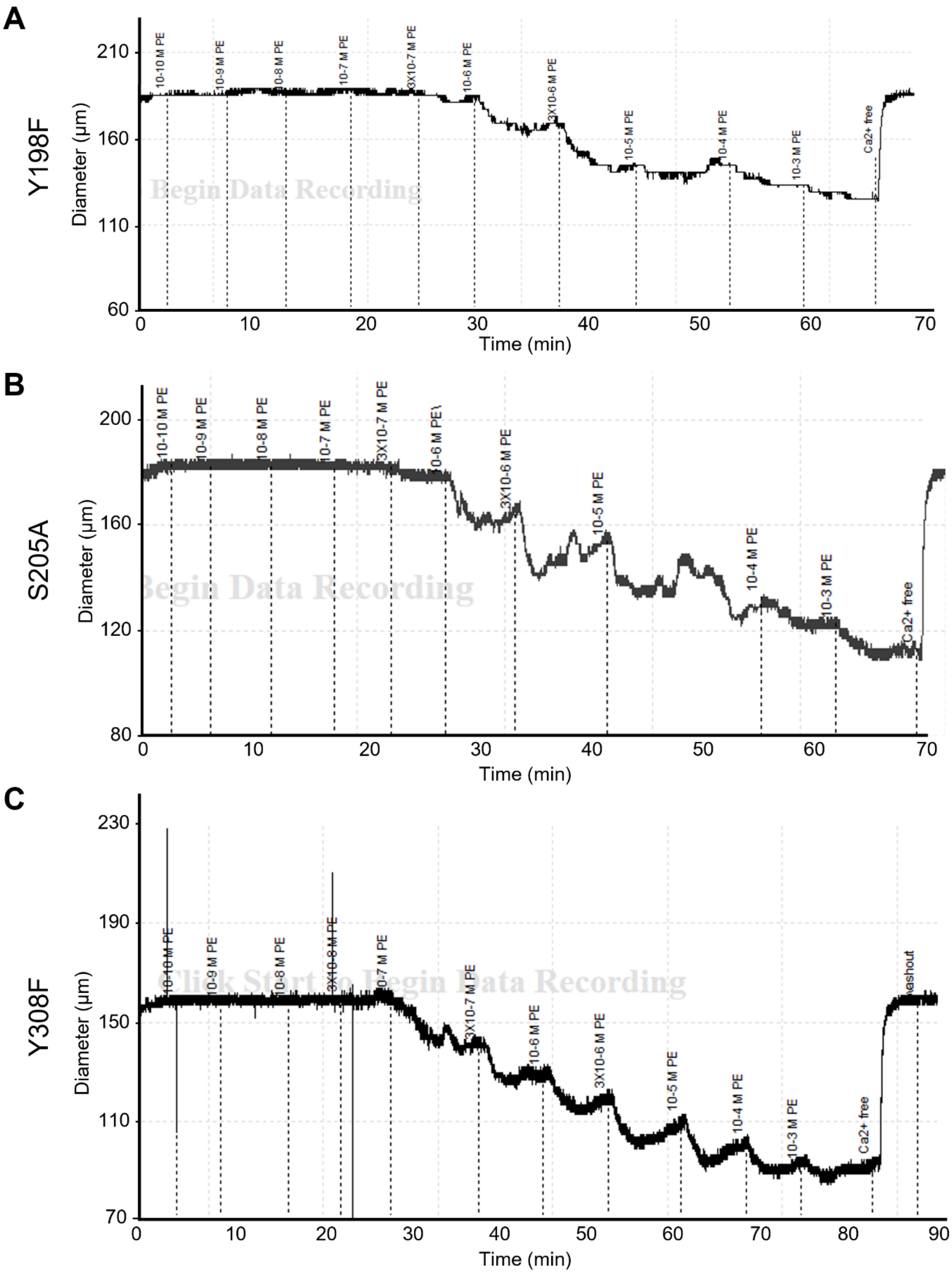


**Fig S4. Representative traces for phenylephrine responses from PANX1 phospho-dead mutant third-order mesenteric arteries.** Pressure myography traces for phenylephrine (PE) dose response curves for Y198F (**A**), S205A (**B**) or Y308F (**C**) third-order mesenteric arteries.

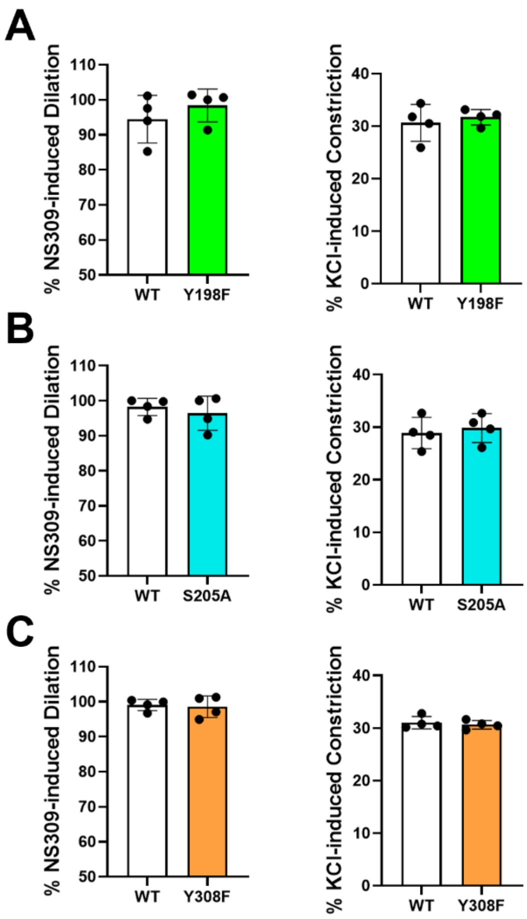


**Fig S5. Endothelial and vascular smooth muscle function in PANX1 phospho-dead mutant mesenteric arteries is intact.** Percent dilation to 1 μM NS309 and % constriction to 30 mM KCl for Y198F (**A**), S205A (**B**) or Y308F (**C**) third-order mesenteric arteries. Unpaired t-test. Bars represent mean ± SD. *N*=5 mice.

**A**

| Gene Symbol | Description | log2 Fold Change | Adjusted *P* Value | Abundance |
|---|---|---|---|---|
| Twist2 | twist family bHLH transcription factor 2 | 2.095180946 | 0.044189365 | Increased |
| Dynlt1b | dynein light chain Tctex-type 1B | 0.699627383 | 0.038641314 | Increased |
| 1700008B11Rik | RIKEN cDNA 1700008B11 gene | -3.885230983 | 0.044189365 | Decreased |

**B**

| Gene Symbol | Description | log2 Fold Change | Adjusted *P* Value | Abundance |
|---|---|---|---|---|
| Arhgap36 | Rho GTPase activating protein 36 | 5.696628758 | 0.005624977 | Increased |
| Gm28932 | Predicted gene 28932 | 5.440978816 | 0.020553628 | Increased |
| Mgarp | Mitochondria localized glutamic acid rich protein | 5.030169815 | 0.042959497 | Increased |
| Spta1 | Spectrin alpha, erythrocytic 1 | 4.177882664 | 0.002552701 | Increased |
| Kl | Klotho | 3.895889516 | 0.044781936 | Increased |
| Kcnh8 | Potassium voltage-gated channel subfamily H member 8 | 3.51772316 | 0.024543007 | Increased |
| Cyp46a1 | Cytochrome P450 family 46 subfamily A member 1 | 2.931377572 | 0.042959497 | Increased |
| Svopl | SVOP like | 2.825320899 | 0.031827899 | Increased |
| Cml3 | N-acetyltransferase 8 family member 3 | 2.63885894 | 0.001159538 | Increased |
| Gm24494 | Predicted gene 24494 | 2.537914871 | 0.029066752 | Increased |
| Slc6a7 | Solute carrier family 6 member 7 | 2.496310545 | 0.027479636 | Increased |
| Cpxm1 | Carboxypeptidase X, M14 family member 1 | 2.462399381 | 4.64E-06 | Increased |
| Nrxn1 | Neurexin 1 | 2.431309755 | 0.002552701 | Increased |
| Adamtsl2 | ADAMTS like 2 | 2.297618896 | 0.027479636 | Increased |
| Tnfaip6 | TNF alpha induced protein 6 | 2.25531241 | 0.002353656 | Increased |
| Pdgfrl | Platelet derived growth factor receptor like | 2.200845544 | 0.024361175 | Increased |
| Adora2a | Adenosine A2A receptor | 2.186989232 | 0.037486627 | Increased |
| Hsd11b1 | Hydroxysteroid 11-beta dehydrogenase 1 | 2.15317643 | 0.037486627 | Increased |
| Epb4.1l3 | Erythrocyte membrane protein band 4.1 like 3 | 2.135933205 | 0.001159538 | Increased |
| Mfap4 | Microfibril associated protein 4 | 2.101972682 | 0.027353394 | Increased |
| Adcy7 | Adenylate cyclase 7 | 2.057032781 | 0.010423055 | Increased |
| Smoc2 | SPARC related modular calcium binding 2 | 2.039545014 | 0.027384168 | Increased |
| Kif1a | Kinesin family member 1A | 2.033943018 | 0.003480825 | Increased |
| Fbln1 | Fibulin 1 | 2.024229676 | 0.024361175 | Increased |
| Gpnmb | Glycoprotein NMB | 2.006675003 | 0.013397034 | Increased |
| Svep1 | Sushi, von Willebrand factor type A, EGF and pentraxin domain containing 1 | 1.996226533 | 0.046912748 | Increased |
| Adcyap1r1 | ADCYAP receptor type I | 1.993175712 | 0.002791244 | Increased |
| Fmod | Fibromodulin | 1.925138961 | 0.016840424 | Increased |
| Wnt5a | Wnt family member 5A | 1.904180761 | 0.024543007 | Increased |
| Sorcs2 | Sortilin related VPS10 domain containing receptor 2 | 1.902996034 | 0.001623699 | Increased |
| Rflna | Refilin A | 1.899247675 | 0.007765512 | Increased |
| Lama2 | Laminin subunit alpha 2 | 1.884626819 | 0.035447112 | Increased |
| Cmtm5 | CKLF like MARVEL transmembrane domain containing 5 | 1.857436115 | 0.030268575 | Increased |
| Gdf10 | Growth differentiation factor 10 | 1.843251109 | 0.035525559 | Increased |
| Slitrk6 | SLIT and NTRK like family member 6 | 1.836402367 | 0.011135073 | Increased |
| Ngfr | Nerve growth factor receptor | 1.82192432 | 0.013220146 | Increased |
| Lum | Lumican | 1.812843297 | 0.048052489 | Increased |
| Ephb2 | EPH receptor B2 | 1.809903675 | 0.002353656 | Increased |
| Itih2 | Inter-alpha-trypsin inhibitor heavy chain 2 | 1.801799566 | 0.024361175 | Increased |
| Gpr153 | G protein-coupled receptor 153 | 1.76871746 | 0.000178788 | Increased |
| Stk32a | Serine/threonine kinase 32A | 1.755243153 | 0.033147574 | Increased |
| Piezo2 | Piezo type mechanosensitive ion channel component 2 | 1.755176266 | 0.010401754 | Increased |
| Cacna1g | Calcium voltage-gated channel subunit alpha1 G | 1.740811368 | 0.001623699 | Increased |
| Shisa3 | Shisa family member 3 | 1.728557566 | 0.024361175 | Increased |
| Lsamp | Limbic system associated membrane protein | 1.696180016 | 0.030268575 | Increased |
| D630003M21Rik | RIKEN cDNA D630003M21 gene | 1.6905143 | 0.002091703 | Increased |
| Cdh11 | Cadherin 11 | 1.681055982 | 0.014877576 | Increased |
| Hunk | Hormonally Increased Neu-associated kinase | 1.679859161 | 0.024038597 | Increased |
| Phlda1 | Pleckstrin homology like domain family A member 1 | 1.678026688 | 0.011135073 | Increased |
| Olfml3 | Olfactomedin like 3 | 1.64944981 | 0.001159538 | Increased |
| Zc3h8 | Zinc finger CCCH type containing 8 | -1.059318093 | 0.049774499 | Decreased |
| Gm44891 | Predicted gene 44891 | -2.780591515 | 0.007852936 | Decreased |
| Gm10699 | Predicted gene 10699 | -3.454187621 | 0.002263472 | Decreased |
| 9130213A22Rik | RIKEN cDNA 9130213A22 gene | -5.148415919 | 0.031462685 | Decreased |

**Fig S6. Top differentially expressed genes in Y198F and S205A mesenteric arteries.** A list of differentially expressed genes (*P*<0.05) from bulk RNA sequencing of Y198F (**A**) and S205A (**B**) mesenteric arteries compared to controls and their relative abundance. Only the top 50 differentially expressed genes with increased abundance are included for S205A mesenteric arteries, ranked by log2 fold change. *N*=3-4 mice.

**A**

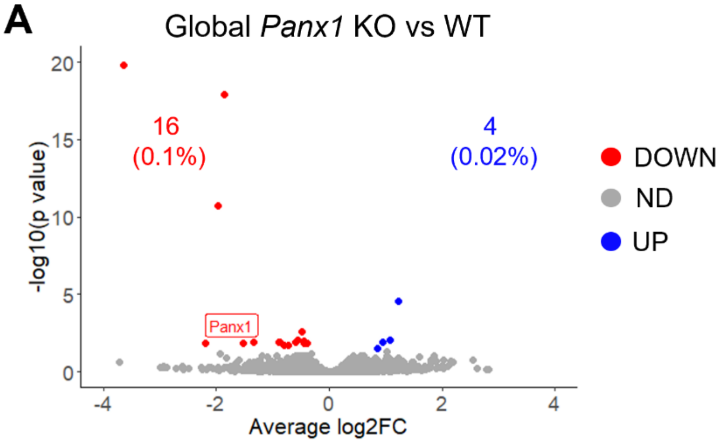


**B**

| Gene Symbol | Description | log2 Fold Change | Adjusted *P* Value | Abundance |
|---|---|---|---|---|
| Dclre1c | DNA cross-link repair 1C | 0.949218238 | 0.012193929 | Increased |
| Chic1 | Cysteine-rich hydrophobic domain 1 | 0.85489228 | 0.030715616 | Increased |
| Pisd-ps1 | Phosphatidylserine decarboxylase pseudogene 1 | 1.080185196 | 0.008853842 | Increased |
| N/A | Novel gene, ENSMUSG00000095041 | 1.226436224 | 2.83E-05 | Increased |
| Adss2 | Adenylosuccinate synthase 2 | -0.490798225 | 0.002392788 | Decreased |
| Gle1 | GLE1 RNA export mediator | -0.452904077 | 0.011183305 | Decreased |
| Rpl7a | Ribosomal protein L7a | -0.382681076 | 0.013286442 | Decreased |
| Ndufs5 | NADH:ubiquinone oxidoreductase core subunit S5 | -0.592704389 | 0.012193929 | Decreased |
| Hrct1 | Histidine-rich carboxyl terminus 1 | -0.898554289 | 0.012193929 | Decreased |
| Gm15446 | Predicted gene 15446 | -3.653433148 | 1.61E-20 | Decreased |
| Chchd2 | Coiled-coil-helix-coiled-coil-helix domain containing 2 | -0.453323174 | 0.014676118 | Decreased |
| Timp4 | Tissue inhibitor of metalloproteinase 4 | -0.871843015 | 0.012193929 | Decreased |
| BC051019 | cDNA sequence BC051019 | -1.340637109 | 0.011647903 | Decreased |
| Gpx4 | Glutathione peroxidase 4 | -0.79274967 | 0.020711986 | Decreased |
| Plac9 | Placenta associated 9 | -1.97174443 | 2.01E-11 | Decreased |
| Tmem254 | Transmembrane protein 254 | -1.852776785 | 1.23E-18 | Decreased |
| Panx1 | Pannexin 1 | -2.185572578 | 0.013286442 | Decreased |
| Cripto | Cripto, EGF-CFC family member | -1.5246884 | 0.014945957 | Decreased |
| Naa38 | N-alpha-acetyltransferase 38 | -0.728513527 | 0.018857579 | Decreased |
| Elob | Elongin B | -0.549012473 | 0.008853842 | Decreased |

**Fig S7. Global *Panx1* KO mesenteric arteries are transcriptomically similar to WT controls.** Bulk RNA sequencing of mesenteric arteries from global *Panx1* knockout (KO) mice compared to wildtype (WT) controls. *N*=3 mice. (**A**) Volcano plot exhibits all differentially expressed genes based on *P*<0.05. Blue dots and text show the number and percentage of genes with increased abundance, red dots and text show the number and percentage of genes with decreased abundance and grey dots represent genes not statistically different (ND) from controls. (**B**) A list of differentially expressed genes in global *Panx1* KO mesenteric arteries compared to WT controls and their relative abundance. The Ensembl ID was included in the description for the novel gene that currently has no gene symbol (N/A).

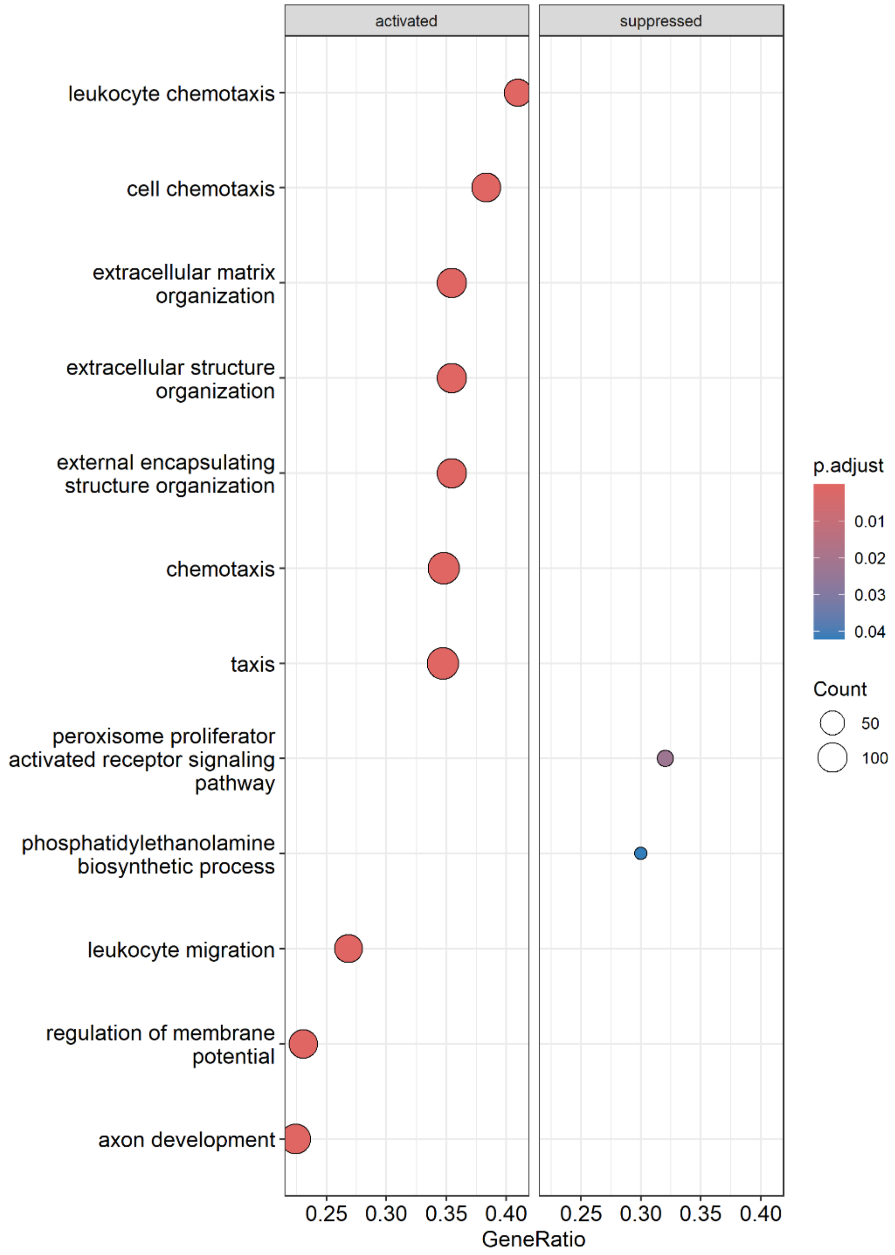


**Fig S8. Membrane potential, migratory and extracellular matrix pathways are activated in S205A mesenteric arteries.** Bulk RNA sequencing of mesenteric arteries from S205A mice compared to littermate controls. Dot plot shows the top ten positively enriched (activated) Gene Set Enrichment Analysis pathways and the only negatively enriched (suppressed) pathways based on an adjusted *P* value (p.adjust) $< 0.05$ and ranked based on normalized enrichment score. Count is the number of core enrichment genes driving the pathway enrichment. GeneRatio is the proportion of pathway genes that contribute to the enrichment score, calculated by count/setSize where setSize is the total number of genes in the pathway. *N*=4 mice.

| Comparison | Protein Interactors |
| --- | --- |
| WT vs Y198F | Pzp, Prdx4, Cdk11b, Ighv1-54, Igkv4-63, Krt7, Pbx3, Klhdc10, Krt4, Pfn1, Brd7, Hoxb9, Atp2a1, Ddx4, Sfxn3, Hspa4, Sptan1, Krt72, Nup133, Tars1, Ddx18, Rbfa, Aldh1l2, Fam133b, Timeless, Jup, Anxa1, Gnao1, Cebpz, Son, Bysl, Smc2, C3, Nr2f2, Inf2, Mndal, Sh2d3c, Actn3, Igkv6-14, Eloc, Vdac3, Slc25a31, Mta3, Ywhaq, Nob1, Itprid2, Tubb1, Serpina1a, Isg20 |
| Y198F vs WT | Ighv6-5, Eif3d, Gm20521, Nop14, Ppil4, Timm50, Emilin1, Atp5pb, Nup205, Sec13, Plekhg3, Dut, Zhx1, Vapa, Ak1, Eif5b, Snw1, Loxl2 |
| WT vs S205A | Msh6, Ighv1-54, Krt7, Pfn1, Nid1, Brd7, Sfxn3, Nsun5, Gm5629, Cdc42 |
| S205A vs WT | Eif3d, Cggbp1, Gpc1, Kifc5b, Plekhg3, Gm20521, Nop14, Rrs1, Ppil4, Ywhae, Kdm1a, Emilin1, Arl1, Eif5b, Unc45a, Lrrfip2, Chchd3, Nup205, Usp39, Acadl, Umps, Acads, Dut, Polr2b, Ccar2, Zhx1, Sart1, Gls, Vapa, Acly, Stip1, Snw1, Por, Loxl2, Timm50 |
| WT vs Y308F | Wdr43, Ilk, Cdk11b, Sf3b4, Igkv4-63, Etfb, Ap2b1, Dcaf13, Pfn1, Rbfox2, Gnas, Tdrd3, Ddx4, Sfxn3, Nsun5, Snrpd2, Phldb2, Fkbp8, Hnrnpll, Srbd1, Aldh1l2, Dkc1, Pdha1, Bclaf1, Nhp2, Mrpl14, Aacs, Cebpz, Pclaf, Igf2bp1, Hells, Trim28, Nsf, Zfp326,Znf326, Ipo4, Ythdf3, Bysl, Ston1, Prpf31, Atxn2, Phf5a, Yars1, Igkv6-14, Ehd1, Bub3, Rnh1, Dhx36, Upf3b, Nob1, Zfp384, Smc3, Anp32b, Cep55 |
| Y308F vs WT | Cggbp1, Pip4p2, Nop14, Ppil4, Emilin1, Bcl7c, Unc45a, Chchd3, Umps, Plekhg3, Kdm1a, Polr2b, Acadl, Zhx1, Arl1, Sart1, Stip1, Loxl2 |
| WT Only | Pfn1, Sfxn3 |
| Y198F Only | Ighv6-5, Atp5pb, Sec13, Ak1 |
| S205A Only | Gpc1, Kifc5b, Rrs1, Ywhae, Lrrfip2, Usp39, Acads, Ccar2, Gls, Acly, Por |
| Y308F Only | Pip4p2, Bcl7c |

**Fig S9. Protein interactor comparisons for PANX1 WT and phospho-dead mutants.** *Panx1* knockout MOVAS were transfected with PANX1 wildtype (WT), Y198F, S205A or Y308F HA-tagged constructs, immunoprecipitated using a mouse HA antibody and sent for mass spectrometry to compare protein-protein interactors. A list of unique protein interactors when comparing PANX1 WT to each phospho-dead mutant and vice versa (vs; where the interactors were detected in the PANX1 variant co-immunoprecipitation listed first) as well as interactors identified in only one PANX1 variant co-immunoprecipitation when comparing to all other variants (Only). *N*=1 co-immunoprecipitation per construct.

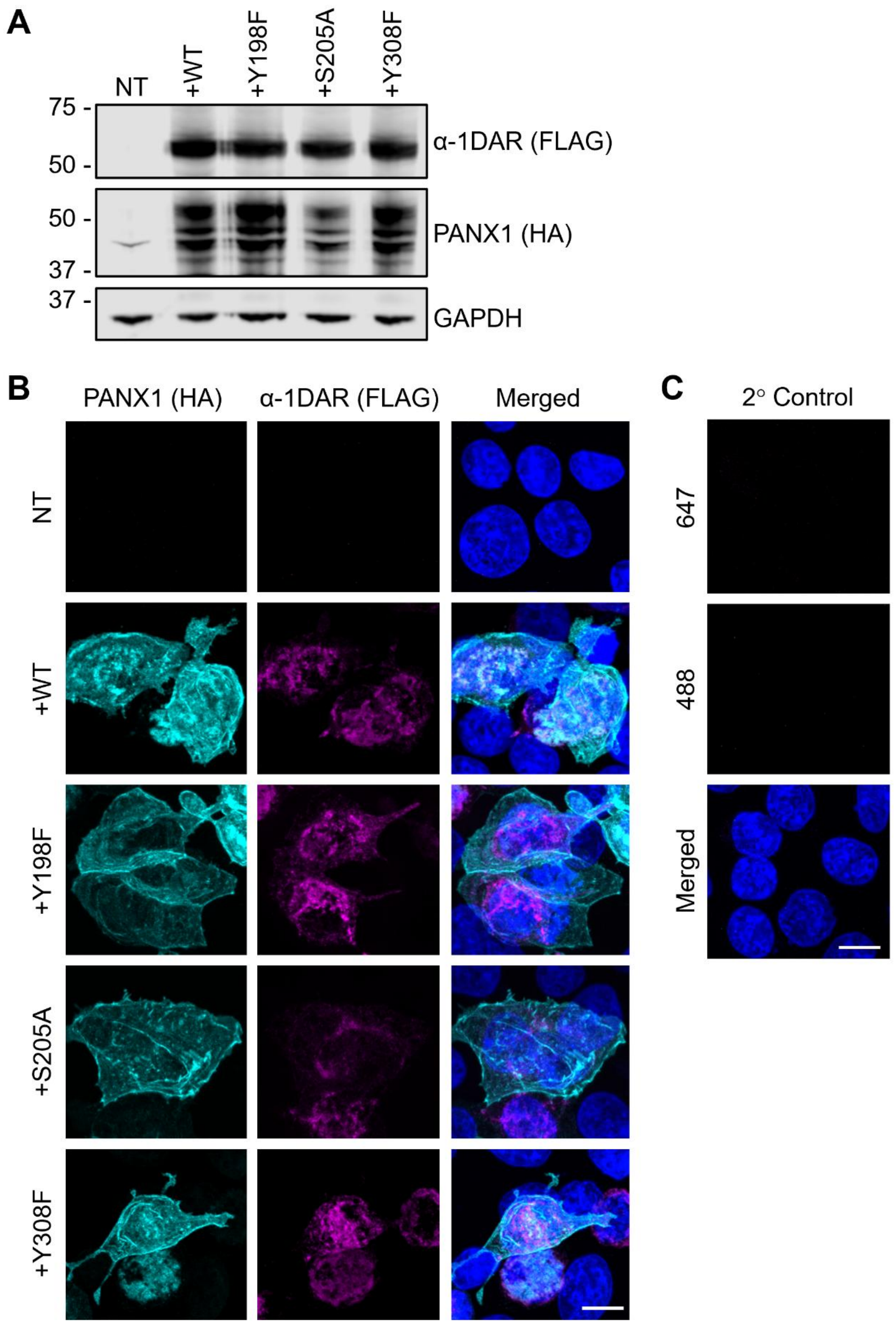


**Fig S10. PANX1 phospho-dead mutants show similar protein levels and localization compared to WT when overexpressed in HEK293T cells.** Human embryonic kidney

(HEK293T) cells were co-transfected with various PANX1 HA-tagged constructs as well as FLAG-tagged α-1D adrenergic receptor (α-1DAR) and assessed by immunoblotting (**A**) and immunocytochemistry (**B**). For the Western blot, GAPDH as protein loading control, protein sizes in kDa. For the confocal micrographs, PANX1 in cyan, α-1DAR in magenta and nuclei stained with DAPI in blue. Scale bar 10 µm. (**C**) Coverslips incubated with Alexa Fluor™ 488 and 647 antibodies as secondary only (2°) control. Scale bar 10 µm. *N*=3 independent transfections. NT, non-transfected; WT, wildtype.

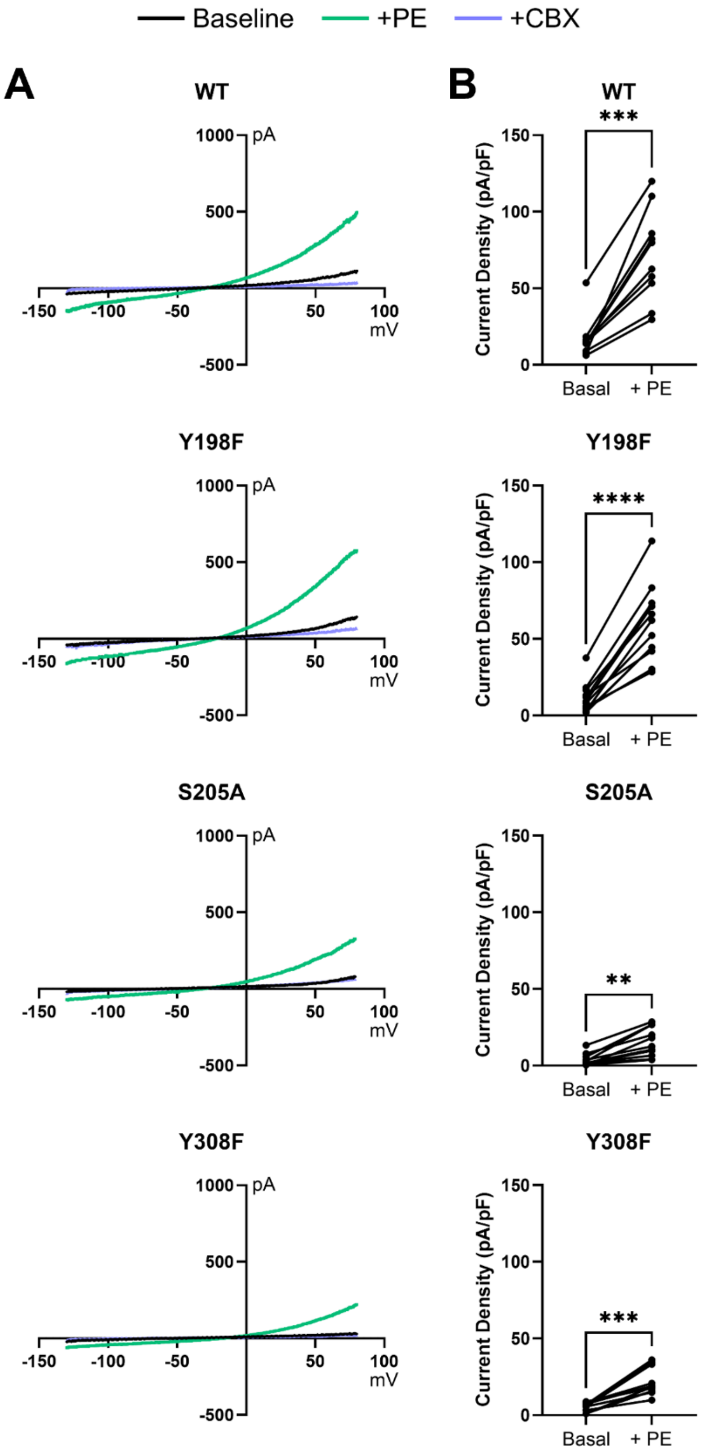


**Fig S11. Current-voltage curves and phenylephrine activation of PANX1 WT and phospho-dead mutant PANX1 channel currents.** (**A**) Representative current-voltage relationship curves from whole-cell patch-clamp recordings of mouse PANX1 wildtype (WT), Y198F, S205A and Y308F in human embryonic kidney (HEK293T) cells co-transfected with the mouse α-1D adrenergic receptor and indicated PANX1 construct at baseline (black line) and with phenylephrine stimulation (+PE, 20 μM; green line). PANX1 currents were inhibited by carbenoxolone (CBX, 100 μM; purple line). (**B**) Current density for PANX1 WT or phospho-dead mutant whole-cell patch-clamp at baseline (basal) and after PE stimulation. Welch's t-test (*n*>18 cells, *N*>5 independent transfections for each construct). ***$P$<0.001, ****$P$<0.0001.

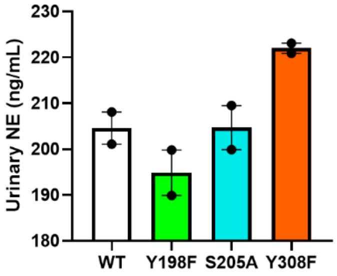


**Fig S12. Urinary norepinephrine levels of PANX1 phospho-dead mutant mice.** Concentrations of norepinephrine (NE) were measured in the urine of wildtype (WT), Y198F, S205A and Y308F mice. *N*=2 mice.

**Data File S1.** Normalized counts, group comparisons and Gene Set Enrichment Analysis of PANX1 phospho-dead mutant and global *Panx1* knockout mouse mesenteric arteries bulk RNA sequencing.

**Data File S2.** PANX1 wildtype and phospho-dead mutant co-immunoprecipitation-mass spectrometry protein cluster and individual protein interactor results and associated EnrichR pathway analyses.
